# Engineering cathepsin S selective chemical probes and antibody-drug conjugates through substrate profiling with unnatural amino acids

**DOI:** 10.64898/2025.12.20.695745

**Authors:** Maria Łęcka, Oliwia Gorzeń, Natalia Ćwilichowska-Puślecka, Julia Nguyen, Martyna Majchrzak, Vanessa Pippa, Bartosz Dołęga-Kozierowski, Piotr Kasprzak, Boris Turk, Marcin Drąg, Rafał Matkowski, Marcin Poręba

## Abstract

Cysteine cathepsins, particularly cathepsin S, are important regulators of proteolytic signaling in health and disease, including cancer progression and immune modulation. Despite their therapeutic relevance, selective chemical tools to study individual cathepsins remain limited due to overlapping substrate preferences. Here we present design of cathepsin S selective chemical probes and cathepsin S-cleavable antibody-drug conjugates (ADCs) through substrate profiling with unnatural amino acids. First, Hybrid Combinatorial Substrate Library (HyCoSuL) technology incorporating a broad spectrum of unnatural amino acids was applied to comprehensively investigate the substrate specificity of cathepsin S. This approach enabled the identification of highly selective tetrapeptide motifs that served as scaffolds for the design of optimized fluorogenic substrates, irreversible inhibitors, and fluorescent activity-based probes (ABPs). These tools demonstrated high selectivity toward cathepsin S over closely related family members in both biochemical and cellular settings. We then translated these findings to develop cathepsin S-activated ADCs, incorporating the optimized peptide motifs as protease-cleavable linkers for targeted payload release. Using these linkers, we further generated MMAE-based antibody-drug conjugates directed against HER-2 and the TROP-2 proteins, demonstrating cathepsin S-dependent cytotoxicity in HER-2-positive as well as HER-2-negative/TROP-2-positive breast cancer models. Finally, we employed anti-cathepsin S antibodies in combination with mass cytometry (CyTOF) to assess the spatial distribution of cathepsin S expression in breast cancer patient samples. By correlating cathepsin S levels with tumor-associated markers such as TROP-2 and HER-2, we identified potential co-expression patterns that support the rationale for personalized therapy using antibody-drug conjugates selectively activated by cathepsin S. This integrative approach provides a comprehensive platform for profiling cathepsin S activity at the molecular, cellular, and tissue levels, with broad implications for the development of precision therapeutics and diagnostic strategies in oncology.

## Introduction

The cysteine protease cathepsin S (CTSS) is a unique member of the cathepsin family, distinguished by its selective expression pattern, functional versatility, and enzymatic stability at neutral to mildly alkaline pH. Under physiological conditions, cathepsin S is predominantly expressed in antigen-presenting cells (APCs), including macrophages, dendritic cells, and B cells, where it plays an important role in adaptive immunity (1, 2). Unlike most cathepsins, cathepsin S remains catalytically active at neutral pH, allowing it to function within lysosomes, cytosol and extracellularly, particularly under inflammatory conditions (3). In addition to its role in antigen processing, this protease has been increasingly implicated in pathological processes, including autoimmune diseases, atherosclerosis, and cancer (4). In the tumor microenvironment (TME), cathepsin S contributes to cancer progression through multiple converging mechanisms: degradation of extracellular matrix (ECM) components, promotion of angiogenesis, and modulation of immune responses (5). Notably, cathepsin S is not only expressed by tumor cells but also secreted by stromal and immune cells, especially tumor-associated macrophages (TAMs), further amplifying its impact within the TME (6, 7). Elevated cathepsin S expression has been reported in various cancers, including prostate, gastric, pancreatic, colorectal, and breast carcinomas, where it is often associated with poor prognosis, increased metastatic potential, and therapeutic resistance (5). In breast cancer, particularly in aggressive subtypes such as triple-negative breast cancer (TNBC) and HER-2-positive tumors, cathepsin S expression is frequently upregulated (8) (9). Given its restricted physiological expression and aberrant activity in cancer, cathepsin S emerges as an attractive target for precision oncology (4). Moreover, its dual function in both immune regulation and tumor biology creates a unique opportunity to exploit CTSS not only as a biomarker for disease progression and patient stratification, but also as a therapeutic target for selective intervention, such as prodrugs activation (10).

The functional versatility of cathepsin S is closely linked to its distinct substrate specificity, which sets it apart from other cathepsins such as B, L, and K. Although cathepsin S shares similarities at the P1 position, cleaving after residues such as Arg, Lys, or Gln, its primary specificity determinant lies in the P2 position (11). Structural and functional studies using natural substrates and combinatorial peptide libraries have shown that the S2 subsite of cathepsin S preferentially accommodates aliphatic and hydrophobic residues such as Leu, Val, Nle, and Met (12). This specificity, along with its ability to function at neutral or even oxidizing pH, contributes to its exceptional proteolytic activity beyond endo-lysosomal compartments (3). For instance, cathepsin S is involved in the cleavage of thyroglobulin in the thyroid follicle lumen, highlighting its compartment-specific function (13). Moreover, degradomics studies have revealed cathepsin S activity at the cell surface, where it acts as a sheddase, cleaving membrane proteins such as cell adhesion molecules, growth factor receptors, and signaling regulators (14). The information about cathepsin S peptide preferences was further translated into chemical probes. Early synthetic substrates for this proteas took advantage of its preference for aliphatic residues at P2 such as Val or Met, yielding fluorogenic and FRET-based probes with that allowed for real-time detection of cathepsin S activity *in vitro* and in antigen-presenting cells (15). These tools showed good selectivity over cathepsins B and L and were used to monitor invariant chain degradation in the MHC class II pathway. To enable *in vivo* imaging, substrate designs evolved to include reverse-engineered constructs, in which non-peptidic cathepsin S inhibitors were modified to include cleavable peptide bonds (16). Lipidated near-infrared fluorescent (NIRF) substrates derived from this strategy showed high selectivity and accumulated at sites of proteolytic activity, such as tumor-associated macrophages, delivering strong fluorescent signals in mouse models. Further refinement led to the development of structurally constrained probes, such as peptide hairpin-based sensors like MG101, which incorporates a cathepsin S-specific cleavage site flanked by an electrostatic zipper and a NIRF dye-quencher pair (17). This design enabled highly selective and pH-tolerant imaging of CTSS activity in macrophages and was validated using cathepsin S knockout tissue. Altogether, synthetic substrates, ranging from soluble peptides to membrane-tethered imaging constructs, have proven valuable for tracking cathepsin S activity with spatial and temporal precision in live cells and animal models. Another tools to track the activtiy of cathepsin S are the inhibitor-like activtiy-based probes (ABPs), that covalently label active enzymes, enabling direct visualization or enrichment from live systems (18). Initial ABPs using epoxysuccinyl electrophiles, such as DCG-04 derivatives, showed limited selectivity and poor retention (19). Subsequent designs, including quenched ABPs (qABPs), offered major improvements (20, 21). For instance, BODIPY-conjugated qABPs enabled real-time *in vivo* imaging of cathepsin S activity in breast tumors, revealing strong, macrophage-associated signals with low background fluorescence (21). More recently, modular two-step ABPs have been developed, incorporating bio-orthogonal handles (azide or alkyne) for post-labeling (22). A cathepsin S-selective two-step probe based on the LHVS scaffold enabled precise subcellular localization of active cathepsin S using confocal and correlative light-electron microscopy. Another major advance came with DOTAM-based activatable probes, which combine cathepsin S-cleavable motifs with targeting ligands such as cRGD peptides (23). These constructs integrate protease activation with tumor-targeting, allowing selective accumulation and activation in cathepsin S- and integrin-positive cancer cells.

Although significant progress has been made in the development of chemical tools targeting cathepsin S, many existing probes still suffer from incomplete selectivity due to overlapping substrate preferences with related cathepsins, and calpains (12, 24). To overcome this limitation, we employed a Hybrid Combinatorial Substrate Library (HyCoSuL) approach to explore the cathepsin S substrate recognition landscape in the P4-P1 positions, incorporating a diverse set of unnatural amino acids (25, 26). By systematically comparing this specificity profile to those of other cathepsins, we identified highly selective peptide motifs, which we used to design optimized substrates, covalent inhibitors, and activity-based probes. Using these ABPs, we evaluated cathepsin S activity in breast cancer models, which guided our selection of triple-negative breast cancer cell lines for subsequent prodrug studies. Recently, we also demonstrated that HyCoSuL-derived peptide linkers containing unnatural amino acids can be successfully repurposed and used to achieve cathepsin B- and L-selective activation in prodrugs and antibody-drug conjugates for cancer therapy (27). Building on this proof-of-concept, the present study applies a similar strategy to cathepsin S, extending the HyCoSuL-guided protease-selective linker approach to this enzyme. This enabled tumor-targeted drug release using the same specificity principles established with our substrates and ABPs. Finally, to support personalized therapeutic strategies, we applied mass cytometry (CyTOF, cytometry by time-of-flight) to map cathepsin S expression in primary breast cancer tissues and correlate its abundance with clinically relevant tumor markers, including canonical receptors such as HER-2 and the emerging ADC target TROP-2. In parallel, we evaluated cathepsin S-selective linkers in ADC formats directed not only against HER-2, but also against TROP-2, to explore how protease-guided linker design can be combined with distinct surface antigens across HER-2-positive and HER-2-negative breast cancer subtypes.

## Material and Methods

### Chemicals, reagents, and antibodies

All chemicals were obtained from commercial suppliers and used without further purification. Fmoc- and Boc-protected amino acids were purchased from Iris Biotech GmbH, Merck Sigma-Aldrich, Combi-Blocks, Angene, Chemat, and Ambeed. The fluorescent dye Fmoc-ACC-OH was obtained from Aapptec Peptides. Peptide and ACC substrate syntheses were performed using Rink amide AM resin (200-300 mesh, loading 0.74 mmol/g) and 2-chlorotrityl chloride resin (2-CTC, 100-200 mesh, loading 1.6 mmol/g), both sourced from Iris Biotech GmbH. Coupling reagents (HATU, HBTU), piperidine, 2,2,2-trifluoroethanol (TFE), and trifluoroacetic acid (TFA) were also from Iris Biotech GmbH. Anhydrous HOBt was purchased from Creosalus. Additional reagents, including 2,4,6-collidine, acetonitrile (ACN), triisopropylsilane (TIPS), 4-aminobenzyl alcohol, and bis(4-nitrophenyl) carbonate, were obtained from Sigma-Aldrich. Solvents such as *N,N′*-dimethylformamide (DMF), methanol (MeOH), dichloromethane (DCM), acetic acid (AcOH), diethyl ether (Et_2_O), and phosphorus pentoxide (P_2_O_5_) were supplied by POCh (Gliwice, Poland). Inhibitors E64, E64d, CA-074 and CA-074Me were purchased from Sigma-Aldrich. All synthesized substrates and peptide prodrugs were purified by reverse-phase high-performance liquid chromatography (RP-HPLC) using a Waters system (M600 solvent delivery module and M2489 detector) equipped with a semi-preparative Discovery® C8 column (10 µm particle size). Compound purity and molecular mass were confirmed by LC-MS (Waters). Trastuzumab (anti-HER-2, A2007) and Sacituzumab (anti-TROP-2, A2031) were obtained from Selleckchem, and monomethylauristatin E (MMAE) was purchased from MedKoo Biosciences.

### Enzyme kinetic studies

All kinetic studies were performed using an fMax fluorescence plate reader (Molecular Devices) in kinetic mode with 96- or 384-well plates. ACC fluorescence was monitored at excitation and emission wavelengths of 355 nm and 460 nm, respectively. Recombinant human cysteine cathepsins L, V, B, K, and S were expressed and purified as previously described (28, 29). Prior to use, all cysteine cathepsins were titrated with E64 to normalize enzymatic activity. Assays for cathepsins were conducted in 100 mM sodium acetate buffer containing 10 mM NaCl and 10 mM DTT (pH 5.5). Breast cancer cell lysates (BT-474, MCF-7, MDA-MB-231) were prepared by culturing cells to confluence in the appropriate media, replacing media with PBS, and harvesting the cells using a cell scraper (without trypsin). The cells were pelleted by centrifugation (400 × g, 5 min), resuspended in PBS, and centrifuged again under the same conditions. After removal of the supernatant, the pellet was resuspended in cathepsin assay buffer (excluding DTT) and sonicated. Lysates were then clarified by centrifugation (12,000 × g, 5 min), and the supernatants were collected and stored at −80 °C until use. All enzymatic assays, including HyCoSuL library screening and kinetic analyses of synthetic substrates, inhibitors, probes and prodrugs, were conducted at 37 °C and repeated at least three times. Mean values are reported. Kinetic data were analyzed using GraphPad Prism software (version 10.4.1).

### Characterization of cathepsin S specificity at the P1 position

To evaluate cathepsin S specificity at the P1 position, a fluorogenic substrate library of the general structure Ac-Ala-Arg-Leu-P1-ACC was used. The library contained 19 natural and over 100 unnatural amino acids. Screening was performed in triplicate at a final substrate concentration of 4 µM and cathepsin S concentration at 5 nM. The total assay time was 30 min; however, only the linear range of substrate hydrolysis (5-15 min) was used for data analysis. The average hydrolysis rate was calculated for each substrate, with standard deviation values below 10%. The substrate Ac-Ala-Arg-Leu-Arg-ACC was used as a reference, with its hydrolysis rate set to 100%, and all other values were normalized accordingly.

### Characterization of cathepsin S specificity at the P4-P2 positions

To investigate cathepsin S specificity at the P4-P2 positions, HyCoSuL libraries with fixed P1-Arg and P1-Gln residues were used. The P4, P3, and P2 sub-libraries of each set were screened at a final concentration of 100 µM in a total volume of 100 µL using human recombinant cathepsin S. The active enzyme concentration ranged from 5 to 25 nM depending on the sub-library. The total assay time was 30 min; however, to avoid substrate depletion, only the linear portion of the reaction (10-15 min) was used for rate calculations (RFU/s). Each screening experiment was performed at least in triplicate, and mean values were used to construct the cathepsin S specificity matrix. The standard deviation for each substrate was below 15%. The hydrolysis rate of the most efficiently cleaved amino acid at each position was set to 100%, and all other values were normalized accordingly. To better contextualize the substrate preferences of cathepsin S, its specificity profile was compared to those of cathepsins B, L, V, and K, which were previously characterized using P1-Arg HyCoSuL libraries by our group (26, 30, 31).

### Fluorogenic substrate synthesis and kinetic analysis

Peptide substrates for cathepsin S were designed based on P4-P1 specificity profiling and synthesized on Rink amide resin using standard Fmoc solid-phase peptide synthesis protocols (32). The resin was swollen in dichloromethane (DCM) and deprotected with 20% piperidine in DMF. Fmoc-ACC-OH was coupled twice using HOBt/DIC in DMF and allowed to react overnight. Subsequent Fmoc-protected amino acids were coupled sequentially using HATU and 2,4,6-collidine, with Fmoc deprotection after each step using 20% piperidine in DMF. Following chain assembly, the N-terminus was acetylated using AcOH/HBTU/DIPEA in DMF for 45 min. Peptides were cleaved from the resin using TFA/TIPS/H_2_O (95:2.5:2.5, v/v/v) for 2 hours, precipitated in cold diethyl ether, lyophilized, and purified by preparative HPLC. Final compounds were dissolved in DMSO at 20 mM and stored at −80 °C. To evaluate substrate selectivity, each fluorogenic peptide (10 µM) was incubated with recombinant cysteine cathepsins (5 nM) in assay buffer at 37 °C. Fluorescence emission was recorded at 460 nm, and initial cleavage rates were reported as relative fluorescence units per second (RFU/s). For selected substrates, detailed kinetic parameters (k_cat_, K_M_, and k_cat_/K_M_) were determined as previously described (33).

### Synthesis of AOMK-based inhibitors

The synthesis of inhibitors and activity-based probes featuring an acyloxymethyl ketone (AOMK) warhead was carried out as previously described (34, 35). In this study, we prepared a panel of cathepsin S inhibitors incorporating six different P1 amino acids: Glu(Me), Lys(2ClZ), Arg, Cys(Bzl), Cys(MeBzl), and Nle(OBzl). The procedure is exemplified with P1-Glu(Me) inhibitor. The synthesis started with the conversion of Boc-Glu(Me)-OH to the corresponding diazomethyl ketone, Boc-Glu(Me)-CH_2_N_2_, using a solution of diazomethane in diethyl ether. This intermediate was subsequently treated with 30% HBr in acetic acid and water (1:2, v/v) to yield Boc-Glu(Me)-CH_2_Br. The resulting crude product (1.0 equiv.), a pale yellow oil, was reacted with 2,6-dimethylbenzoic acid (2,6-DMBA, 1.2 equiv.) in the presence of potassium fluoride (KF, 3.0 equiv.) in DMF to obtain Boc-Glu(Me)-AOMK. After deprotection with 50% trifluoroacetic acid (TFA) in dichloromethane (DCM), the free acid (Glu(Me)-AOMK) was used directly in the coupling step without further purification. Other P1-AOMK fragments were synthesized analogously. In parallel, selected P4-P3-P2 peptide fragments bearing appropriate protecting groups were synthesized on 2-chlorotrityl chloride resin and cleaved under mild conditions. Two peptide fragments, Ac-Met(O_2_)-Cit-NptGly and Ac-Phe(F_5_)-Cit-Lys(2ClZ)-OH, were obtained and used without additional purification. For the final coupling step, the peptide fragments (1.0 equiv.) were reacted with their corresponding P1-AOMK moieties (1.2 equiv.) in DMF using HATU and DIPEA (1.2 equiv. each) as coupling agents. The crude products were purified by reverse-phase high-performance liquid chromatography (HPLC), lyophilized, and reconstituted in DMSO to a final concentration of 10 mM.

### Synthesis of fluorescent activity-based probes

Fluorescently labeled activity-based probes (ABPs) targeting cathepsin S were synthesized using a protocol analogous to that described for the corresponding unlabeled inhibitors. Initially, the peptide precursor Boc-PEG(4)-Phe(F_5_)-Cit-Lys(2ClZ)-COOH was synthesized on 2-chlorotrityl chloride resin and used without further purification. In parallel, the warhead Boc-Glu(Me)-AOMK was synthesized as previously outlined. Following Boc deprotection of the warhead using a 50% TFA/DCM mixture, the resulting Glu(Me)-AOMK (1.0 equiv.) was coupled to Boc-PEG(4)-Phe(F_5_)-Cit-Lys(2ClZ)-COOH (1.3 equiv.) in DMF using HATU/DIPEA (1.3 equiv. each) to yield Boc-PEG(4)-Phe(F_5_)-Cit-Lys(2ClZ)-Glu(Me)-AOMK. The crude product was purified by reverse-phase HPLC, after which the Boc group was removed with TFA in DCM. Residual TFA was removed under a stream of argon. The resulting free amine (1.0 equiv.) was subsequently labeled with either Cy5-NHS or BODIPY-NHS (1.2 equiv.) in DMF to afford the final fluorescent probes: Cy5-PEG(4)-Phe(F_5_)-Cit-Lys(2ClZ)-Glu(Me)-AOMK and BODIPY-PEG(4)-Phe(F_5_)-Cit-Lys(2ClZ)-Glu(Me)-AOMK. The labeled probes were purified via HPLC, analyzed by LC-MS, and dissolved in DMSO at a final concentration of 10 mM. Using the same synthetic strategy, two additional ABPs were generated: Cy5-PEG(4)-Met(O_2_)-Cit-NptGly-Glu(Me)-AOMK and BODIPY-PEG(4)-Met(O_2_)-Cit-NptGly-Glu(Me)-AOMK.

### Kinetic analysis (k_obs_/[I]) of inhibitors and activity-based probes

The second-order rate constants of inhibition (k_obs_/[I]) were determined for human recombinant cathepsins S, B, L, and V using their respective assay buffers. All measurements were performed under pseudo-first-order kinetic conditions, as previously described (34). Inhibitors or ABPs were serially diluted in assay buffer and transferred to a 96-well plate. Each well was supplemented with the fluorogenic substrate Cbz-Phe-Arg-AMC at a final concentration of 25 µM and preincubated for 15 minutes. In parallel, enzymes were pre-activated in assay buffer for 15 minutes at room temperature. Following preincubation, the activated enzymes were added to the wells, and fluorescence was monitored immediately for 30 minutes. The k_obs_/[I] values (M^−1^S^−1^) were calculated from the progress curves and averaged from at least three independent experiments. Final inhibitor (or ABP) concentrations were maintained at a minimum of fivefold excess relative to enzyme concentrations to ensure pseudo-first-order conditions. Results were analyzed in GraphPad Prism and are reported as mean ± standard deviation.

### SDS-PAGE-based profiling of cathepsin reactivity with OG-233 ABP

To assess the potency and selectivity of the OG-233 activity-based probe, active site-titrated human recombinant cathepsins were individually preincubated at a final concentration of 10 nM in assay buffer for 15 minutes at 37 °C. Subsequently, each enzyme preparation was incubated with OG-233 at final probe concentrations of 10 nM, 100 nM, and 500 nM for 30 minutes in a total reaction volume of 200 µL. After incubation, 100 µL of 3× SDS loading buffer containing DTT was added to each sample. The mixtures were boiled for 5 minutes, and 20 µL of each sample was loaded onto 4-12% Bis-Tris Plus gels (15-well format). Electrophoresis was performed at 200 V for 30 minutes alongside 2 µL of PageRuler™ Prestained Protein Ladder. Following separation, the gels were scanned directly at 700 nm (Cy5 fluorescence channel, excitation 685 nm) using a Sapphire™ biomolecular imager (Azure Biosystems). Band intensities corresponding to labeled cathepsins were quantified using Image Studio software.

### Detection of active cathepsin S in breast cancer cell using Cy5-labeled OG-233 and OG-234 probes

50,000 of MDA-MB-231 cells were seeded into 12-well plate and allowed to attach overnight. The next day, OG-233 and OG-234 probes (1 μM final concentration) were added to the cells and incubated for various times (from 0,5 h up to 24 h). Control cells were preincubated with cathepsin S inhibitors (JN1, JN4, JN7) at 25 μM for 2 hours, and then incubated with OG-233 or OG-234 probes for 24 hours. Next, cells were harvested and prepared for SDS-PAGE analysis as described in the above section. Electrophoresis was run for 25 min (4-12% Bis-Tris Plus 10-well gel, 200 V). Next, proteins were transferred onto the membrane (0.2 μm nitrocellulose membrane, 10 V, 60 min) and Ponceau S was used to verify equal loading and transfer. Membranes were then blocked with 5% BSA in TBS-T buffer for 1 hour at RT. Membranes were next incubated anti-cathepsin S antibody (1:1,000, overnight, 4 °C), and the next day with a secondary antibody (IRDye® 800CW, donkey anti-Goat) for 30 min at RT, and scanned at 680 nm and 800 nm using the Sapphire™ biomolecular imager (Azure Biosystems) to detect the Cy5 probes and cathepsin S. All images were analyzed with Image Studio software.

### Synthesis of peptide-based prodrugs

Tetrapeptide prodrugs (OG-313, OG-314 and OG-352) were synthesized via standard Fmoc solid-phase peptide synthesis (SPPS) on 2-chlorotrityl chloride (2-CTC) resin using 5 mL polyethylene syringe reactors. Sequential Fmoc deprotection and amino acid coupling cycles were performed to construct the P4-P3-P2-P1 sequence on resin. Following completion of chain assembly, the N-terminus was acetylated using acetic acid, HBTU, and DIPEA in DMF. Peptides were cleaved from the resin using a DCM/TFE/AcOH mixture (8:1:1, v/v/v), and residual solvents were removed by evaporation with hexane. The crude peptides were dissolved in acetonitrile/water (3:1, v/v) and lyophilized. The resulting Ac-tetrapeptides, bearing a free carboxylic acid at the C-terminus, were conjugated to *p*-aminobenzyl alcohol (PABOH, 1.3 equiv.) using HATU (1.0 equiv.) and 2,4,6-collidine (2.0 equiv.) in DMF for 30 minutes to 3 hours at room temperature. Reaction progress was monitored by LC-MS. Crude Ac-tetrapeptide-PABOH intermediates were purified by preparative HPLC, lyophilized, and confirmed by LC-MS. To attach the self-immolative linker, Ac-tetrapeptide-PABOH (1.0 equiv.) was reacted with bis(4-nitrophenyl) carbonate (BPNPC, 2.0 equiv.) and DIPEA (2.0 equiv.) in DMF for 24 hours. If necessary, 0.006 equiv. of HOBt was added to facilitate the reaction. The resulting Ac-tetrapeptide-PABC-PNP intermediate was purified by HPLC and characterized by LC-MS. In the final step, Ac-tetrapeptide-PABC-PNP (1.0 equiv.) was coupled to monomethyl auristatin E (MMAE, 1.2 equiv.) in the presence of DIPEA (2.0 equiv.) in DMF at room temperature for 4 hours. Completion of the reaction was confirmed by LC-MS. The final product, Ac-tetrapeptide-PABC-MMAE, was purified by preparative HPLC, lyophilized, and verified by LC-MS. Purified prodrugs were dissolved in DMSO at a final concentration of 20 mM for storage and further use.

### Kinetic analysis of prodrug cleavage by LC-MS

Cleavage assays were conducted at 37 °C in 1 mL reaction volumes using acetate buffer composed of 100 mM sodium acetate (pH 5.5), 100 mM NaCl, 1 mM EDTA, and 5 mM DTT. Buffers were prepared at room temperature and equilibrated before use. Prodrugs were initially dissolved in DMSO at a stock concentration of 10 mM, then diluted into the reaction buffer to a final concentration of 50 µM, maintaining a DMSO concentration below 0.5% (v/v). Reactions were initiated by the addition of recombinant cathepsin and carried out in glass LC-MS vials incubated at 37 °C. At specified time points (0, 5, 30, 60, and 120 minutes), aliquots were taken and directly injected into the LC-MS system without further processing. Quantification of MMAE release was based on absorbance at 220 nm. The area under the MMAE peak was integrated, and the amount of released MMAE was plotted as a function of time. Each time point was measured in triplicate, and results are presented as mean ± standard deviation (SD). A standard curve was prepared using free MMAE at concentrations of 1, 5, 10, 20, and 50 µM to enable accurate quantification of MMAE released from the prodrugs.

### Cell culture

The BT-474 (ATCC HTB-20, RRID: CVCL_0179), MCF-7 (ATCC HTB-22, RRID: CVCL_0031), and MDA-MB-231 (ATCC HTB-26, RRID: CVCL_0062) breast cancer cell lines were cultured at 37 °C in a humidified incubator with 5% CO_2_. Cells were maintained in Dulbecco’s Modified Eagle Medium (DMEM; Gibco, Cat. No. 41965-039) supplemented with 10% fetal bovine serum (FBS; Gibco, Cat. No. A5256801), L-glutamine, sodium pyruvate, penicillin (100 U/L), and streptomycin (0.1 mg/mL). All cell lines were routinely tested for mycoplasma contamination (every 12-16 weeks) using a PCR-based detection kit.

### Detection of cathepsin S in breast cancer cell lines

Human breast cancer cell lines BT-474, MCF-7, and MDA-MB-231 were harvested and lysed directly in reducing SDS sample buffer. Lysates were denatured by heating at 95 °C for 5 minutes and resolved by SDS-PAGE on 4-12 % Bis-Tris gels at 200 V for 25 minutes. Proteins were transferred to 0.2 µm nitrocellulose membranes using wet transfer at 10 V for 60 minutes. Following transfer, membranes were blocked in 5 % bovine serum albumin (BSA) in TBS-T (Tris-buffered saline with 0.1% Tween-20) for 1 hour at room temperature. Membranes were then incubated overnight at 4 °C with a goat anti-human cathepsin S primary antibody (Bio-Techne, AF1183) diluted 1:1,000 in TBS-T containing 1% BSA. After three washes (10 minutes each) in TBS-T, membranes were incubated for 30 minutes at room temperature with an Alexa Fluor 790-conjugated rabbit anti-goat IgG (H+L) secondary antibody (ThermoFisher, A27019) at a 1:10,000 dilution. Membranes were then washed three additional times in TBS-T and imaged at 790 nm using the Sapphire™ biomolecular imager (Azure Biosystems).

### Synthesis of ADCs and their quality control following conjugation

Synthesis of ADCs. Trastuzumab or sacituzumab (200 μg) were dissolved in 100 μL of PBS (pH 7.4) and incubated with a 10-fold molar excess of TCEP for 30 min at 37°C to reduce interchain disulfide bonds and generate free thiols. A 10-fold molar excess of maleimide-peptide-PABC-MMAE (in DMSO) was then added, and the reaction mixture was incubated for an additional 30 min at 37°C. Unreacted peptide-drug was removed using 30 kDa MWCO Amicon centrifugal filters (12,000 × g, 8 min, RT). The retentate (ADC) was washed once with PBS and centrifuged again under the same conditions (12,000 × g, 8 min, RT). ADC concentration was determined by absorbance at 280 nm and adjusted to 1 mg/mL. ADCs were stored at 4°C until use. To assess ADC integrity, ADCs and trastuzumab (1 μg/sample) were analyzed by SDS-PAGE. To assess the structural integrity of antibody-drug conjugates (ADCs), both ADC samples and unconjugated trastuzumab or sacituzumab (1 µg per sample) were analyzed under reducing and non-reducing conditions. Samples were mixed with SDS loading buffer (with or without reducing agent), heated at 95 °C for 5 minutes, and resolved by SDS-PAGE on 4-12% Bis-Tris gels at 200 V for 25 minutes. Protein bands were visualized by overnight staining at room temperature using InstantBlue™ Coomassie Protein Stain (Abcam, Cat. No. ab119211). Gels were imaged at 650 nm using the Sapphire™ biomolecular imager (Azure Biosystems) to assess the purity and molecular weight distribution of ADCs compared to the parent antibody.

### TROP-2 internalization asaay

MDA-MB-231 cells were seeded at a density of 20,000 cells/mL onto poly-L-lysine-coated microscope slides and incubated overnight. The following day, cells were washed and incubated for 30 min with goat anti-TROP-2 antibody (R&D Systems, AF650) diluted 1:50 in culture medium. Cells were then fixed, and bound anti-TROP-2 antibody was detected using an Alexa Fluor 488-conjugated mouse anti-goat secondary antibody (*vendor, cat #*). After washing, nuclei were counterstained with Hoechst dye (1:1,000) for 5 min at room temperature. Following additional washes, cells were imaged using a Leica DMi8 fluorescence microscope (Leica Microsystems, Wetzlar, Germany).

### MTS cell viability assay

Cell viability was evaluated using the MTS assay (CellTiter 96® AQueous One Solution Cell Proliferation Assay, Promega, Cat. No. G358B) according to the manufacturer’s instructions. BT-474 cells were seeded at a density of 20,000 cells/well, while MCF-7 and MDA-MB-231 cells were seeded at 5,000 cells/well in 96-well plates and allowed to adhere overnight. Peptide prodrugs, free MMAE (used as a control), or ADCs were added in serial dilutions. For peptide prodrugs and MMAE, concentrations ranged from 0.001 to 10 µM; for ADCs, from 0.03 to 3 µM. After a 6-hour incubation period, the treatment medium was removed and replaced with fresh culture medium. Cells were then incubated for an additional 96 hours (4 days). Following the incubation period, MTS reagent was added directly to each well, and plates were incubated at 37 °C for 3 hours. Absorbance was measured at 490 nm. Background absorbance (blank wells) was subtracted from all readings, and cell viability was calculated as a percentage relative to untreated controls. Data are presented as mean ± standard deviation (SD) from at least three independent experiments unless otherwise stated.

### Synthesis of a panel of metal-labeled antibodies

All antibodies were conjugated to metal-chelating polymers using the MaxPar® X8 Antibody Labeling Kit (Standard BioTools) following the manufacturer’s protocol (36). Briefly, each purified antibody was first buffer-exchanged into the provided R buffer, reduced with TCEP, and then incubated with a lanthanide-loaded polymer to generate stable metal-labeled conjugates. For the tumor architecture and immune phenotype panel, the following metal-conjugated antibodies were prepared: anti-cPARP (^143^Nd), anti-cleaved caspase-3 (^142^Ce), anti-CD66b (^113^Cd), anti-CD45 (^89^Y), anti-CD19 (^165^Ho), anti-CD3 (^154^Sm), anti-CD4 (^110^Cd), anti-CD8 (^114^Cd), anti-CD11b (^209^Bi), anti-CD56 (^176^Yb), anti-EpCAM (^141^Pr), anti-Cadherin-3 (^172^Yb), anti-FAP (^164^Dy), anti-SMA (^161^Dy), and anti-CD31 (^144^Nd). In addition, to investigate the relationship between cathepsin S expression and classical breast cancer markers, we labeled antibodies against HER-2 (^113^Cd),TROP-2 (^168^Er), progesterone receptor (PR, ^145^Nd), estrogen receptor (ER, ^163^Dy), cathepsin B (^162^Dy), and cathepsin S (^155^Gd).

### Single cell CyTOF analysis of tumor samples

Fresh tumor specimens were obtained from four female patients with grade 2 breast cancer under a protocol approved by the Bioethical Commission at Wroclaw Medical University (KB-135/2021) and the Institutional Review Board of the Lower Silesian Oncology, Pulmonology and Hematology Center. Patients were enrolled between November 2021 and December 2022 and ranged in age from 45 to 87 years. Blinding and power analysis were not applicable to this exploratory study. All patients were diagnosed and treated at the Department of Surgical Oncology, Breast Unit, Wroclaw Medical University – Lower Silesian Oncology, Pulmonology and Hematology Center, Wroclaw, Poland. The study was conducted in accordance with the Declaration of Helsinki, and all participants provided written informed consent prior to sample collection. To analyze cathepsin S expression across different breast cancer subtypes, the patient cohort included tumors with diverse receptor profiles, including ER-, PR-, and HER-2-positive as well as triple-negative cases. Patients with suspected distant metastases, a history of other malignancies, lack of written consent, or current pregnancy/lactation (in women of childbearing potential) were excluded. Fresh tumor samples (∼50-250 mm^3^ fragments) were processed within 2 hours of surgical resection. Tissues were transferred to gentleMACS™ C Tubes containing 5 mL of pre-warmed enzyme mix from the Human Tumor Dissociation Kit (Miltenyi Biotec) and mechanically dissociated using the MACS® Octo Dissociator. The resulting cell suspensions were filtered through a 70 µm nylon mesh, washed with PBS (300 × g, 5 min), and pelleted. Red blood cells were lysed using 2 mL RBC lysis buffer for 5 minutes at room temperature, followed by two PBS washes. Viable cells were counted using trypan blue exclusion and adjusted to a concentration of 1-3 × 10^6^ cells/mL for downstream CyTOF staining. Cells were first incubated with a panel of in-house metal-conjugated antibodies targeting surface markers: CD66b, CD45, CD11b, CD19, CD3, CD4, CD8, CD56, CD31, HER-2, PR, ER, TROP2, Cadherin-3, EpCAM, FAP, and SMA. Incubation was carried out at room temperature for 30 minutes. After staining, cells were washed three times in Maxpar® Cell Staining Buffer (CSB), fixed in 1.6% paraformaldehyde for 10 minutes, and permeabilized using Perm-S buffer (Thermo Scientific). Permeabilized cells were washed once with PBS and incubated for 2 hours at room temperature with metal-conjugated anti-cathepsin S antibody in Perm-S buffer. After intracellular staining, cells were centrifuged (900 × g, 5 min), washed once with PBS, then once with Perm-S buffer. Cells were subsequently stained with the Ir191/Ir193 DNA intercalator for 30 minutes at room temperature. Following nuclear staining, cells were centrifuged again (900 × g, 5 min), resuspended in 500 µL CyFACS buffer, and stored at 4 °C for up to 12 hours prior to CyTOF acquisition. Before acquisition, cells were washed twice with CyFACS buffer and twice with Cell Acquisition Solution (Standard BioTools). Mass cytometry data were analyzed using Cytobank (RRID: SCR_014043), employing viSNE-CUDA and FlowSOM algorithms (RRID: SCR_016899). Results are presented as viSNE plots (37). A detailed protocol for immune cell identification using mass cytometry, including staining, sample acquisition, data normalization, and analysis, was previously described by Bagwell et al. (38).

## Results and Discussion

### Cathepsin S substrate specificity at P4-P1 positions

Cathepsin S is a lysosomal cysteine protease (papain-like family) distinguished by its selective expression profile and unique functional features antigen-presenting cells. It plays a critical role in MHC class II antigen processing via degradation of the invariant chain (2, 39). Unlike most other cathepsins, cathepsin S remains proteolytically active at neutral pH, enabling it to function in both endolysosomal and extracellular environments (3, 40). This unusual pH stability is accompanied by a substrate-recognition profile that is both broad and finely tuned, setting cathepsin S apart from close homologs such as cathepsins B, L, V, and K. Early studies with positional scanning combinatorial libraries (PS-SCLs) and individual substrates showed that cathepsin S imposes its most stringent specificity at the P2 position (2, 12). Only a limited set of amino acids, primarily aliphatic and branched side chains such as valine, leucine, norleucine, and methionine, are efficiently tolerated at this position. In contrast, polar or charged residues at P2 significantly impair substrate hydrolysis. This marked P2 selectivity distinguishes cathepsin S from related enzymes such as cathepsin L, which prefers aromatic residues like phenylalanine at the same site. Meanwhile, the S3 and S4 pockets of cathepsin S are more permissive, accommodating a wide variety of amino acids, although acidic residues such as aspartate are generally disfavored. At the P1 position, cathepsin S shows a preference for arginine and lysine, consistent with other cysteine cathepsins, but it also efficiently recognizes glutamine, threonine, and methionine. While these findings highlighted some distinguishing features of cathepsin S specificity, particularly at the P2 position, they also underscored the considerable overlap with other family members, making it difficult to achieve true selectivity using only natural amino acids (12). To explore the chemical space of the cathepsin S active site more comprehensively and to identify structural motifs that could improve selectivity, we employed the Hybrid Combinatorial Substrate Library approach incorporating a wide range of unnatural amino acids (25). This method has previously been used to profile other cathepsins, including B, L, and K, yielding detailed specificity profiles that have guided the development of selective chemical probes and inhibitors (26, 30, 31). We first profiled the P1 specificity of cathepsin S using a fluorogenic Ac-Ala-Arg-Leu-P1-ACC substrate library containing 19 natural and over 100 unnatural amino acids. As expected, natural residues such as Arg, Lys, Gln, and Thr were well tolerated. However, several unnatural residues, such as Cys(Bzl), Cys(MeBzl), Nle(OBzl), Lys(2ClZ), and Glu(Bzl), were hydrolyzed with significantly greater efficiency than the best natural amino acids (**Figure 1**). This revealed that the P1 pocket of cathepsin S is not only permissive but can strongly accommodate bulky, hydrophobic, and chemically modified side chains. Notably, although many of the top P1 amino acids were also accepted by other cathepsins, several residues, such as Glu(Me), Cit, Lys(tfa), and Thr(Bzl), demonstrated promising differential activity that could aid in designing cathepsin S-selective substrates (**Figure S1**). Next, we examined substrate recognition at the P4-P2 positions using a HyCoSuL library with fixed P1-Arg (**Figure 2, Figure S2**). Cathepsin S showed a clear preference at P2 for aliphatic residues such as dhAbu, 2Aoc, NptGly, Leu, and Nle, with additional recognition of structures like 4Pal, Lys(2ClZ), and Ala(2Th). When these results were compared with corresponding specificity data for cathepsins B, L, V, and K, several striking differences emerged. In particular, residues like dhAbu and 2Aoc were more favorably cleaved by cathepsin S than by the other cathepsins. P3 selectivity was narrower, with strong preferences for Phg, Idc, and Pip. Although these were also tolerated by other cathepsins, less commonly recognized residues such as Cit and Thz may offer opportunities for enhancing selectivity. The P4 site, as expected, was relatively promiscuous, accommodating a range of amino acids but showing the highest activity for Met(O_2_). To assess whether P4-P2 specificity was dependent on the residue in the P1 position, we repeated the screen using a P1-Gln HyCoSuL library (**Figure S3**) (41). The observed preferences were largely consistent with those obtained from the P1-Arg library, confirming that P4-P2 recognition is relatively independent of P1 identity in this context. In summary, our comprehensive profiling of cathepsin S substrate specificity across P4-P1 positions confirms many previously known features for natural amino acids but, more importantly, identifies several unnatural amino acids that are cleaved with superior efficiency and selectivity. These findings open the door to the development of highly selective chemical tools, including substrates, activity-based probes, and inhibitors, specifically tailored to cathepsin S.

**Figure 1.**
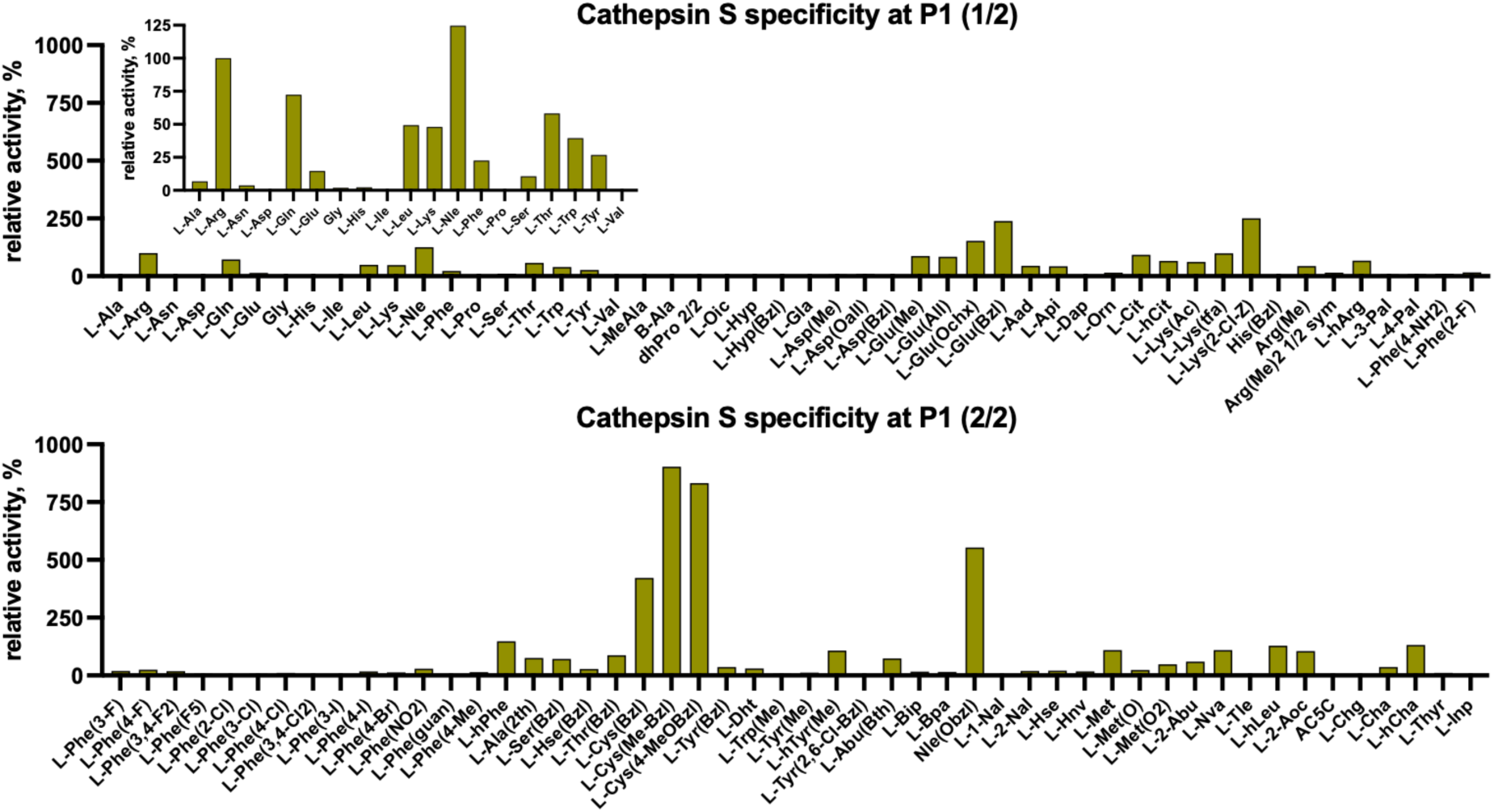
Human cathepsin S specificity at the P1 position. The P1 substrate preference of human cathepsin S was determined using a fluorogenic substrate library based on the Ac-Ala-Arg-Leu-P1-ACC scaffold, containing 19 natural and over 100 unnatural amino acids. The x-axis displays abbreviated amino acid names, while the y-axis shows the relative enzymatic activity for each substrate, normalized to the Ac-Ala-Arg-Leu-Arg-ACC control substrate (set to 100%). The specificity screening was performed in triplicate. Substrate hydrolysis rates (RFU/s) are presented as average values, with standard deviation (SD) below 10% for each substrate. The most efficiently recognized P1 residues were Cys(MeBzl), Cys(Me)Bzl, and Nle(OBzl). As several unnatural amino acids were significantly better recognized than natural ones, a separate graph displaying P1 specificity toward natural amino acids (normalized to Arg = 100%) is included for improved visualization.

**Figure 2.**
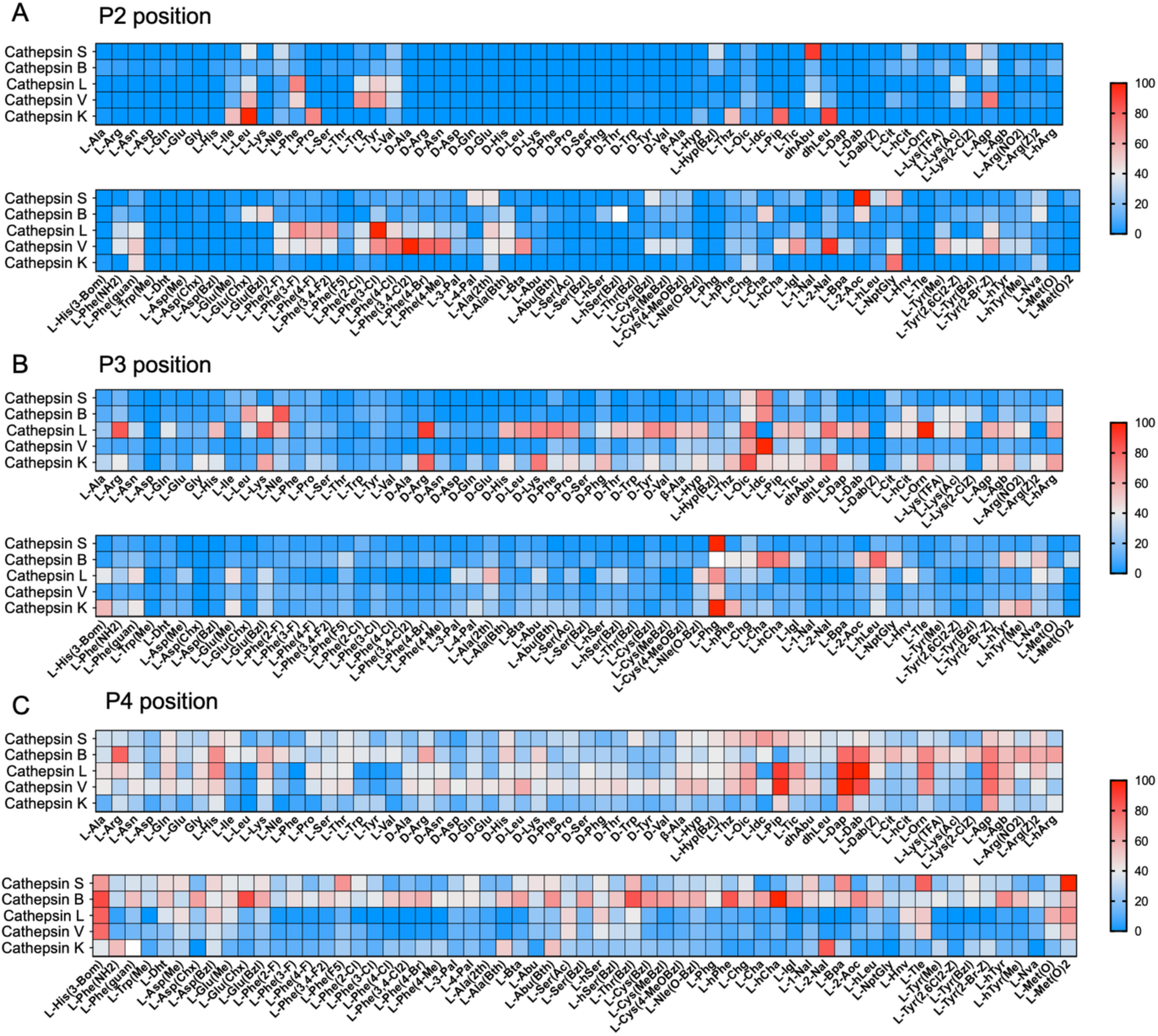
Human cathepsin S specificity at the P4-P2 positions. P4-P2 substrate preferences of human cathepsin S were determined using a HyCoSuL fluorogenic substrate library with fixed P1-Arg. The x-axis displays abbreviated amino acid names, and the y-axis shows the relative enzymatic activity normalized to the best-recognized amino acid at each position: 2Aoc at P2, Phg at P3, and Met(O_2_) at P4 (highlighted in red). Each sub-library (P4, P3, P2) was screened in triplicate, and substrate hydrolysis data (RFU/s, %) are presented as mean values. Standard deviation (SD) was below 10% for each substrate. Cathepsin S specificity profiles were compared with those of cathepsins B, L, V, and K, which were previously characterized using the same P1-Arg HyCoSuL platform (as reported in prior publications). For each enzyme, the most preferred amino acid at a given position is marked in red, and the relative activities of other residues are visualized accordingly using a heat map.

### The development of cathepsin S selective substrates

Since cathepsin S plays key roles in antigen presentation and tumor progression, it represents a valuable target for functional activity assays. Early efforts utilized generic fluorogenic substrates such as Z-Leu-Arg-MCA or Ac-Val-Val-Arg-AMC, but these lacked specificity toward cathepsin S (11). The development of internally quenched FRET substrates (e.g., Abz-LEQ-EDDnp or Mca-GRWPPMGLPWE-K(Dnp)) enabled more selective detection of cathepsin S activity and have since been applied in both biochemical and cellular contexts, including live imaging (15, 42). These substrates vary in their fluorophore-quencher systems, ranging from AMC/AFC to IQF or near-infrared dyes, but their performance depends primarily on the central peptide sequence. Regardless of the detection platform, it is the peptide motif that dictates selectivity and determines the utility of a substrate across applications. Although many substrate architectures are available, highly selective peptide sequences capable of clearly distinguishing cathepsin S from closely related proteases remain limited. Notably, Hu et al. applied a reverse design strategy that converted a potent cathepsin S inhibitor into a fluorescent substrate by introducing a cleavage site near the catalytic center (16). This lipidated probe showed improved selectivity and enabled in vivo imaging of cathepsin S activity in tumors. However, the broader applicability of such substrates across different platforms and biological systems remains constrained by their structural complexity and limited tunability. To address this gap, we conducted a comprehensive substrate specificity screen of cathepsin S and directly compared it with the preferences of other cysteine cathepsins. In the first step, we synthesized a panel of 42 fluorogenic substrates (1st generation) with the general structure Ac-P4-P3-P2-Arg-ACC, where the P4-P2 positions were randomized with both natural and unnatural amino acids, and P1 was fixed as Arg (**Figure 3A, Table S1**). Screening this library revealed several sequences that were efficiently cleaved by cathepsin S and, importantly, showed minimal activity toward cathepsins B, L, and V. The most active substrate was Ac-Met(O_2_)-Cit-2Aoc-Arg-ACC, while Ac-Phe(F_5_)-Cit-2Aoc-Arg-ACC demonstrated the best selectivity. Using this latter sequence as a scaffold, we next synthesized a second-generation library of 16 substrates with variable P1 residues (**Figure 3B, Table S2**). This screen identified Glu(Me) as the most selective P1 residue, retaining strong cathepsin S activity. As expected, Arg, Gln, and Cit were also efficiently cleaved and showed moderate selectivity. Surprisingly, several substrates containing bulky hydrophobic P1 residues were poorly hydrolyzed by cathepsin S. Further analysis revealed that these peptides either precipitated at the screening concentration ([S] = 10 µM) or exhibited significant substrate cooperativity, which hindered efficient binding to the active site. To overcome this, we prioritized more hydrophilic P1 residues in subsequent designs. The third-generation library was constructed based on the most favorable amino acids identified in earlier screens, with the goal of fine-tuning combinations across P4 to P1 (**Figure 3C, Table S3**). Detailed kinetic characterization (k_cat_, K_M_, and k_cat_/K_M_) revealed that several substrates had remarkably low K_M_ values (below 1μM) and, more importantly, high selectivity for cathepsin S (**Table S4, S5, S6**). Two lead substrates, **OG-209** (Ac-Met(O_2_)-Cit-NptGly-Glu(Me)-ACC) and **OG-197** (Ac-Phe(F_5_)-Cit-Lys(2ClZ)-Glu(Me)-ACC), demonstrated over 50-fold selectivity (based on k_cat_/K_M_) compared to cathepsins B and L (**Figure 3D**). This is particularly significant given that cathepsin S generally exhibits lower turnover rates for short peptide substrates compared to its homologs (11, 43, 44). Through this iterative approach incorporating unnatural amino acids, we successfully mapped key cooperativity patterns within the cathepsin S active site and developed a set of highly selective peptide substrates.

**Figure 3.**
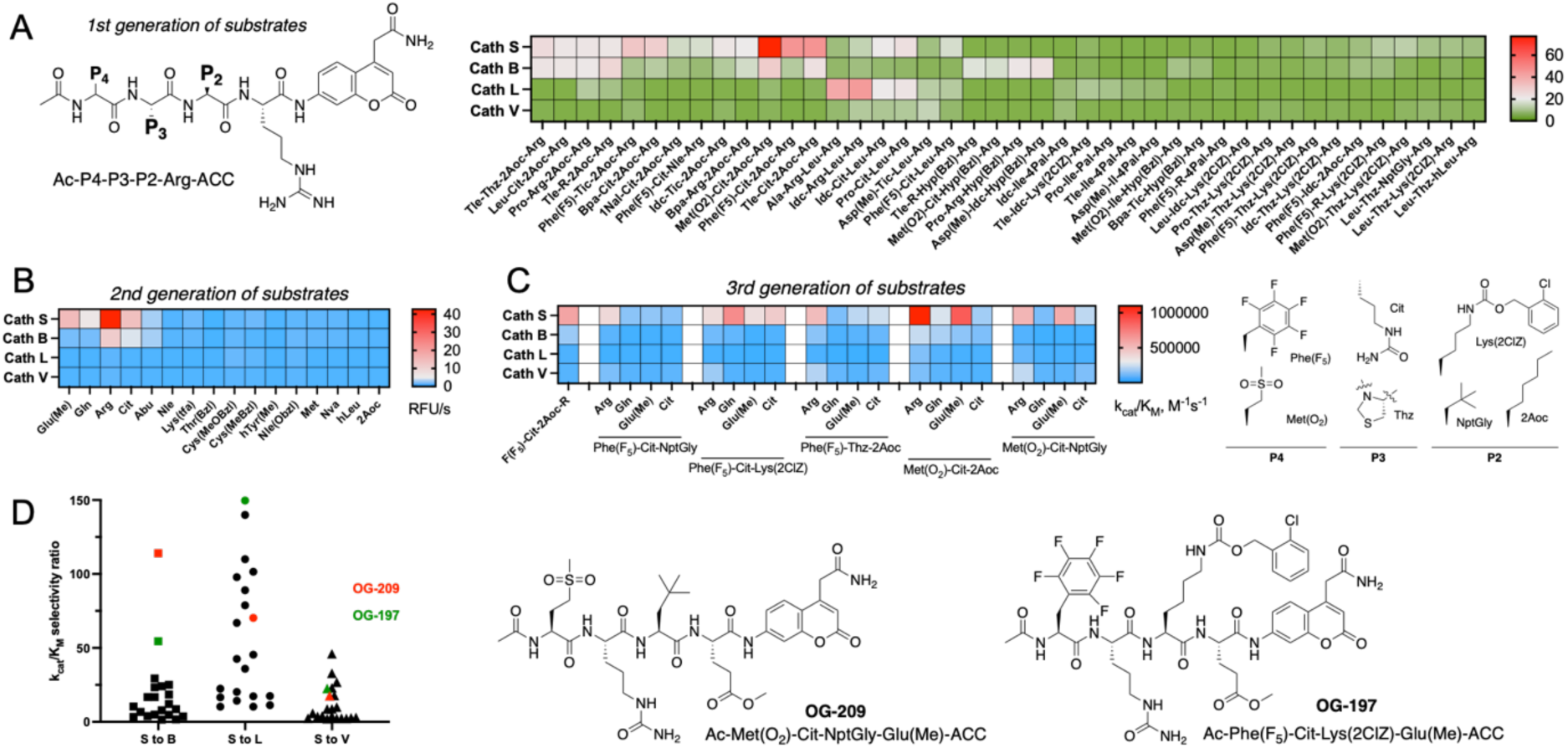
Development of cathepsin S-selective substrates using unnatural amino acids. **A** General structure of the first-generation Ac-P4-P3-P2-Arg-ACC substrates, in which the P4-P2 positions were systematically varied using unnatural amino acids selected based on HyCoSuL screening for cathepsin S. These substrates were tested against cathepsins S, B, L, and V. The hydrolysis rates normalized to enzyme concentration (RFU/s/nM) were used to generate a specificity matrix visualized as a heat map (red: highest activity; green: no cleavage). **B** Specificity matrix generated for second-generation Ac-Phe(F_5_)-Cit-2Aoc-P1-ACC substrates to identify the most selective P1 amino acids for cathepsin S. **C** Specificity profile of third-generation cathepsin S-selective substrates, based on catalytic efficiency (k_cat_/K_M_, M^−1^s^−1^), with optimized P4-P2 residues and variable P1 amino acids. The chemical structures of the most preferred P4-P2 amino acids are shown on the right. **D** Selectivity of third-generation substrates, presented as the ratio of k_cat_/K_M_ for cathepsin S relative to other cathepsins. The structures of the two most selective and highly active substrates, ***OG-209*** and ***OG-197***, are shown on the right-hand side.

### The development of cathepsin S selective inhibitors and activity-based probes

To monitor cathepsin S activity and validate it as a functional target, researchers have designed peptide-based covalent inhibitors and ABPs that exploit the enzyme’s active-site cysteine. Covalent inhibitors typically rely on electrophilic warheads (e.g., vinyl sulfones, nitriles, or acyloxymethyl ketones) that irreversibly acylate or alkylate the catalytic residue, effectively silencing cathepsin S function (4). One classic example is the dipeptidyl vinyl sulfone LHVS, an irreversible cathepsin S inhibitor with nanomolar potency that has served as both a tool compound and structural template (45). Another approach involves the development of non-peptidic nitrile-based inhibitors through substrate activity screening (SAS), which enabled the identification of highly selective cathepsin S inhibitors with favorable potency and reduced cross-reactivity toward other cathepsins (46). ABPs expand upon this concept by coupling the inhibitor scaffold with a reporter tag, such as a fluorophore, quencher pair, or heavy metal isotope, to enable visualization or enrichment of the active enzyme (21, 22). When bound covalently to cathepsin S, the ABP provides a direct readout of enzyme activity, circumventing the limitations of transcript or protein-level measurements, which cannot distinguish between active and inactive forms. For example, Oresic Bender *et al.* developed a quenched fluorescent ABP (BVM157) using a peptide-acyloxymethyl ketone scaffold that remains nonfluorescent until it binds cathepsin S (21). This probe demonstrated high *in vivo* selectivity, enabling clear imaging of this protease activity in tumor-bearing mice. Despite the availability of diverse ABP architectures, from fluorescent to near-infrared and even radiolabeled formats, their selectivity and efficacy are primarily determined by the peptidic recognition element. Therefore, the key to enhancing probe performance lies in the design of peptide sequences that can distinguish cathepsin S from closely related proteases. Building upon this principle, we used a substrate-guided design approach incorporating unnatural amino acids to develop potent and selective cathepsin S-targeting inhibitors and ABPs. We first synthesized a panel of AOMK-based covalent inhibitors using peptide scaffolds derived from our two most selective substrate sequences. These included Ac-Met(O_2_)-Cit-NptGly-P1-AOMK and Ac-Phe(F_5_)-Cit-Lys(2ClZ)-P1-AOMK, in which the P1 position was varied with either Arg or selected unnatural amino acids (**Figure 4A, Table S7**). Kinetic analyses (k_obs_/[I]) revealed that, in both scaffolds, incorporating Glu(Me) at P1 yielded the highest selectivity for cathepsin S over cathepsins B and L (**Figure 4B, Table S8**). Other unnatural residues, such as Cys(Bzl) and Nle(OBzl), also provided substantial selectivity, thereby expanding the repertoire of usable P1 motifs depending on the biological context. Next, we converted these inhibitors into fluorescent activity-based probes (ABPs) by N-terminal labeling with Cy5 or BODIPY (**Figure 4C, Table S9**). Given that bulky fluorophores can influence both potency and selectivity, we carried out detailed kinetic profiling. Remarkably, probes bearing the Phe(F_5_)-Cit-Lys(2ClZ)-Glu(Me) sequence, namely **OG-233** (Cy5) and **OG-235** (BODIPY), displayed excellent selectivity for cathepsin S over cathepsin B (>150-fold), cathepsin L (>2000-fold), and cathepsin V (>500-fold), while also exhibiting over twofold higher potency than the unlabeled parent inhibitor (**Figure 4D**). In contrast, probes based on the Met(O_2_)-Cit-NptGly-Glu(Me) scaffold showed reduced potency and selectivity, reflecting trends observed in the corresponding substrate data. Next, the selectivity and potency of the Cy5-labeled **OG-233** probe was confirmed via SDS-PAGE and fluorescent gel scanning, where increasing concentrations of ABP labeled recombinant cathepsin S over other cathepsins (**Figure 4E**). Finally, we tested whether these ABPs can detect cathepsin S activtiy in living cells. We selected triple negative breast cancer cell line MDA-MB-231. Since these cells belong to immunologically cold tumors, the overall expression of cathepsin S is very low. The neglibile amount of cathepsin S was indeed verified by the antibody. Nevertheless, Cy5-labelled OG-233 and OG-234 probes were able to detect active cathepsin S as soon as upon 3 hours incubation (**Figure 4F**). However, the probe labeled also cathepsin B, and this labeling was far more pronounced than for cathepsin S. Having in mind, that both probes are much more potent toward cathepsin S in kinetic assays, our observation proves the overwhelming expression and activity of cathepsin B. Interestingly, the pretreatment of cells with cathepsin S inhibitors (JN1 or JN7) did not influence probe binding. Importantly, the ABP experiments allowed us to functionally assess cathepsin S activity in triple-negative MDA-MB-231 cells and to identify this line as a suitable model in which active cathepsin S is present at detectable levels. This biological validation created an essential link between our peptide-optimization efforts and their potential translational application, ensuring that the same selective motifs could be repurposed as cleavable linkers in therapeutic designs. Thus, the insights gained from specificity profiling and ABP labeling directly influenced/guided the subsequent development of cathepsin S-cleavable peptide prodrugs and ADCs.

**Figure 4.**
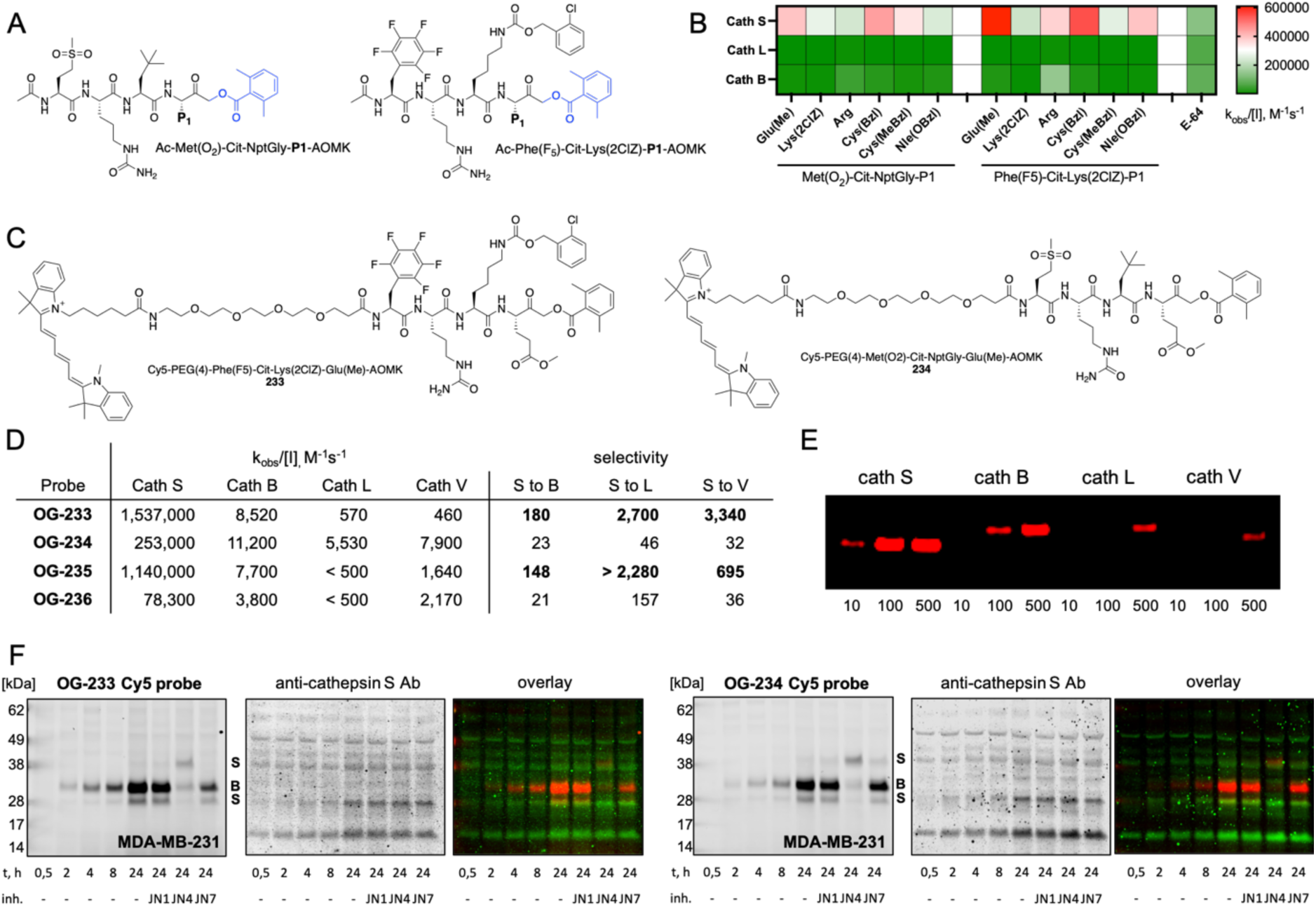
Development of cathepsin S-selective inhibitors and activity-based probes (ABPs). **A** Structures of two series of cathepsin S inhibitors, Ac-Met(O_2_)-Cit-NptGly-P1-AOMK and Ac-Phe(F_5_)-Cit-Lys(2ClZ)-P1-AOMK, designed based on the most selective substrate scaffolds to identify optimal P1 residues for inhibitor potency and selectivity. **B** Specificity of cathepsin S inhibitors toward other cysteine cathepsins (B and L), presented as heat maps of second-order rate constants (k_obs_/[I], M-^1^s^−1^). Red indicates highly potent inhibitors; green indicates weak or inactive compounds. **C** Structures of Cy5-labeled cathepsin S-selective activity-based probes containing Glu(Me) at the P1 position, derived from the most selective inhibitors. **D** Kinetic analysis of Cy5- and BODIPY-labeled probes, including k_obs_/[I] values for cathepsin S and calculated selectivity over cathepsins B, L, and V based on these values. **E** SDS-PAGE analysis of cathepsin labeling by Cy5-labeled OG-233 ABP. Active cathepsins (10 nM) were incubated with increasing probe concentrations (10, 100, and 500 nM), followed by fluorescent gel scanning to visualize probe binding. **F** The labeling of active cathepsin S in MDA-MB-231 live cells with Cy5-tagged OG-233 and OG-234 probes (red). ABPs (1 μM) were incubated with cells for various times from 0,5 h to 24 h with or without cathepsin S inhibitors (JN1, JN4 or JN7). Total cathepsin S was detected with anti-cathepsin S antibody (green).

### Analysis of cathepsin-S cleavable peptide prodrugs and antibody-drug conjugates

Protease-activated prodrugs and peptide-linked antibody-drug conjugates are designed to remain inactive in systemic circulation and become selectively activated within the tumor microenvironment, where protease activity is often upregulated (10, 47). This approach is especially attractive for treating triple-negative breast cancer, a highly invasive and aggressive subtype characterized by elevated protease levels (7, 48). However, most peptide linkers currently used in prodrugs, such as the widely adopted Val-Cit, lack specificity and are cleaved by multiple cathepsins, but more importantly by non-cathepsin proteases, leading to off-target activation and increased systemic toxicity (49, 50). To overcome this limitation, there is a growing focus on developing linkers based on highly selective peptide sequences tailored to tumor-associated proteases (10). Cathepsin S represents a particularly compelling protease target for this strategy. Unlike most other cathepsins, cathepsin S remains catalytically active at neutral pH, allowing it to function not only in endolysosomal compartments but also in the extracellular tumor milieu (17). In tumors, this protease is frequently secreted by both cancer cells and infiltrating immune cells, particularly macrophages and dendritic cells, and retains its activity after secretion (5). This unique extracellular activity enables cathepsin S to activate diffusible prodrugs in the tumor microenvironment, potentially bypassing the need for full cellular internalization. Moreover, cathepsin S is typically restricted to antigen-presenting cells in healthy tissues but becomes upregulated in several malignancies, including TNBC. These features position this enzyme as a promising target for selective prodrug and ADC activation in aggressive cancers. To develop cathepsin S-selective prodrugs, we used optimized peptide sequences derived from our HyCoSuL substrate screen and conjugated them to the potent cytotoxin monomethyl auristatin E (MMAE) via a self-immolative para-aminobenzyl carbamate (PABC) spacer. MMAE is a highly cytotoxic microtubule-disrupting agent that inhibits tubulin polymerization, arresting the cell cycle at the G2/M phase and ultimately inducing apoptosis (51). Due to its high toxicity, MMAE is unsuitable for systemic administration as a free drug and must be delivered in an inactive, masked form. Peptide-MMAE prodrugs remain non-toxic until enzymatically cleaved, at which point the peptide is removed, PABC undergoes self-immolation, and the free drug is released. In our previous work we developed cathepsin B-and cathepsin L-selective prodrugs by using HyCoSuL-derived selective peptides containing unnatural amino acids (27). In this study, we applied the same approach for cathepsin S. We synthesized two cathepsin S-cleavable prodrugs: **313** (Ac-Phe(F_5_)-Cit-Lys(2ClZ)-Glu(Me)-PABC-MMAE) and **314** (Ac-Met(O_2_)-Cit-NptGly-Glu(Me)-PABC-MMAE), along with a non-cleavable control, **313D**, containing DGlu(Me) at the P1 position, and the broadly cleaved pan-cathepsin prodrug Cbz-Val-Cit-PABC-MMAE, **300** (**Figure 5A, Table S10**). To evaluate enzymatic selectivity, we performed LC-MS-based kinetic analysis by monitoring free MMAE release over time in the presence of recombinant cathepsins (**Figure 5B**). As expected, Val-Cit-linked prodrugs were efficiently cleaved by cathepsins S, B, L, and V. In contrast, **313** was selectively and efficiently hydrolyzed only by cathepsin S, demonstrating successful translation from substrate to prodrug. **313D** remained stable, confirming the necessity of an L-amino acid at P1 for efficient cleavage. **314** was also cleaved by cathepsin S, though with moderate cross-reactivity toward other cathepsins, suggesting a lower degree of selectivity than **313**. To assess biological activity, we tested these prodrugs in three breast cancer cell lines: TNBC-derived MDA-MB-231, and HER-2-positive luminal lines BT-474 and MCF-7. Clinical data show that cathepsin S is expressed in TNBC and largely absent in ER-positive subtypes (8), which we confirmed via immunoblotting, detecting cathepsin S only in MDA-MB-231 cells (**Figure 5C**). The presence of active cathepsin S in this cell line was previously confirmed using our ABPs (**Figure 4F**). Our prior studies established that MMAE prodrugs readily enter cells, allowing us to evaluate cytotoxicity following a 6-hour treatment and 4-day incubation to capture the delayed effects of mitotic arrest (27). In MDA-MB-231 cells, the cathepsin S-selective prodrug **313** (Ac-Phe(F_5_)-Cit-Lys(2ClZ)-Glu(Me)-PABC-MMAE) showed potent, concentration-dependent cytotoxicity with an EC_50_ of 27 nM (**Figure 5D**). By contrast, the non-cleavable control **313D** with DGlu(Me) at the P1 position displayed minimal activity (EC_50_ > 1,000 nM), confirming the necessity of proteolytic activation. The second cathepsin S-targeted prodrug **314** and the pan-cathepsin **300** Z-Val-Cit-PABC-MMAE were also active (EC_50_ = 56 nM and 233 nM, respectively), although their reduced potency may reflect less efficient cellular uptake. Next, we tested the same compounds in BT-474 cells, which lack cathepsin S expression. BT-474 cells are inherently more sensitive to MMAE, as shown by the high potency of **300** Z-Val-Cit-PABC-MMAE (EC_50_ = 31 nM). If **313** were broadly cleaved by multiple cathepsins, we would expect a similar or lower EC_50_ in this cell line. Instead, **313** exhibited reduced potency (EC_50_ = 187 nM), indicating that its activity depends primarily on cathepsin S. As expected, the non-cleavable **313D** showed the weakest toxicity (EC_50_ = 575 nM). To further validate selectivity, we generated antibody-drug conjugates by coupling the cathepsin S-cleavable linker to trastuzumab (targeting HER-2) and sacituzumab (targeting TROP-2) (**Figure 5E**). We synthesized cathepsin S-selective ADCs (313 and 314 sequences), a pan-cathepsin Val-Cit ADC (300), and a non-cleavable control ADCs (313D and 314D sequences). None of the trastuzumab-containing conjugates exhibited toxicity in MDA-MB-231 cells, which lack HER-2 expression (**Figure 5F**). However, in HER-2-positive, and cathepsin S-negative BT-474 cells, the pan-cathepsin Val-Cit ADC (300) displayed strong cytotoxicity. In contrast, cathepsin S-selective **313** ADC and the non-cleavable **313D** ADC was only weakly active, comparable to trastuzumab alone, suggesting that its cytotoxicity stems from antibody targeting, not MMAE release. As triple-negative breast cancer lacking HER-2 expression remains a major clinical challenge, we evaluated the cytotoxic activity of cathepsin S-cleavable ADCs incorporating sacituzumab to target the TROP-2 glycoprotein. In MDA-MB-231 cells, the anti-TROP-2 antibody bound specifically to cell-surface TROP-2 and underwent rapid endosomal/lysosomal internalization (**Figure 5G**). We then performed cytotoxicity assays with cathepsin-cleavable and non-cleavable ADC variants. The **313** anti-TROP-2 ADC exhibited moderate toxicity, whereas its non-cleavable counterpart **313D** was almost inactive. In contrast, the **314** anti-TROP-2 ADC was markedly more potent, while the corresponding non-cleavable **314D** showed minimal cytotoxicity (**Figure 5H**). The difference in cytotoxic potency between **313** and **314** likely reflects divergent selectivity toward individual cathepsins. Although both linker peptides are rapidly cleaved by cathepsin S, LC-MS analysis revealed that upon prolonged incubation they are also processed by cathepsins B and L, which are highly abundant in MDA-MB-231 cells. This broader protease susceptibility may accelerate MMAE release and thereby enhance the overall cytotoxic effect of the 314 ADC. Together, these findings demonstrate that cathepsin S-selective peptide sequences derived from HyCoSuL profiling can be successfully translated into protease-cleavable prodrugs and ADCs. Notably, the sacituzumab-based TROP-2 ADCs extend this concept to HER-2-negative/TROP-2-positive TNBC, indicating that cathepsin S-selective linkers can be flexibly combined with alternative antigens when HER-2 is absent. Importantly, these tools target cathepsin S-expressing cancer cells while remaining largely inert in cathepsin S-negative contexts. However, further *in vivo* validation and pharmacological optimization will be essential to fully assess their therapeutic potential in anticancer applications.

**Figure 5.**
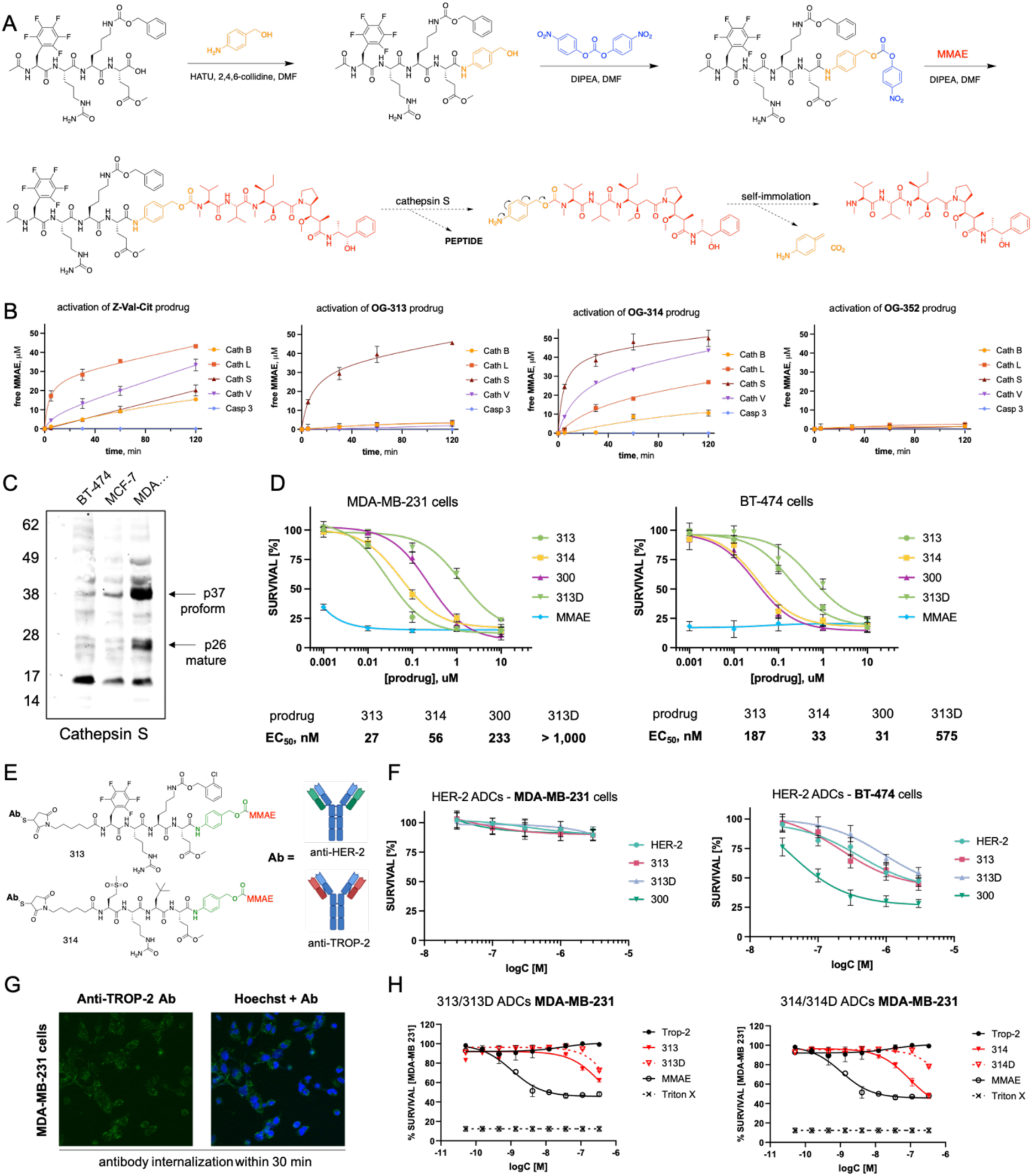
Cathepsin S-cleavable peptide prodrugs and antibody-drug conjugates (ADCs). **A** Schematic representation of the synthesis of cathepsin S-cleavable peptide prodrugs with the general structure Ac-peptide-PABC-MMAE, where PABC (orange) is a self-immolative linker and MMAE (red) is the cytotoxic payload monomethyl auristatin E. The lower panel (dashed arrows) illustrates the mechanism of enzymatic cleavage and subsequent release of free MMAE. **B** Kinetics of MMAE release from two cathepsin S-cleavable prodrugs, OG-313 (Ac-Phe(F_5_)-Cit-Lys(2ClZ)-Glu(Me)-PABC-MMAE) and OG-314 (Ac-Met(O_2_)-Cit-NptGly-PABC-MMAE), compared to a pan-cathepsin-cleavable control (Cbz-Val-Cit-PABC-MMAE) and a non-cleavable control OG-352 (Ac-Phe(F_5_)-Cit-Lys(2ClZ)-_D_Glu(Me)-PABC-MMAE) with a _D_-amino acid at the P1 position. Cleavage and MMAE release by various cathepsins and caspase-3 (negative control) were quantified by LC-MS. **C** Immunoblot analysis of cathepsin S expression in breast cancer cell lines, showing moderate expression in MDA-MB-231 (triple-negative) and absence of expression in BT-474 (HER-2⁺) and MCF-7 (ER⁺/PR⁺) cells. **D** Cytotoxicity of the pan-cathepsin prodrug (Cbz-Val-Cit-PABC-MMAE) and cathepsin S-selective peptide prodrugs in MDA-MB-231 and BT-474 cell lines. The y-axis represents cell viability (%), and the x-axis shows ADC concentration on a log10 scale, ranging from 1 nM to 10 µM. Cell-permeable free MMAE was included as a control. **E** Schematic illustration of the antibody-drug conjugate (ADC) design, exemplified by an anti-HER-2 and anti-TROP-2 ADC incorporating a cathepsin S-cleavable peptide linker (313 Phe(F_5_)-Cit-Lys(2ClZ)-Glu(Me) or 314 Met(O2)-Cit-NptGly-Glu(Me)), a PABC self-immolative spacer, and an MMAE payload. **F** Cytotoxicity of the anti-HER-2 ADCs in HER-2⁺ BT-474 and triple-negative MDA-MB-231 cells. The y-axis represents cell viability (%), and the x-axis shows ADC concentration on a log10 scale, ranging from 0.03 to 3 µM. **G** The internalization of anti-TROP-2 antibody into HER-2 negative MDA-MB-231 cell line performed by fluorescence microscopy. **H** Cytotoxicity of the anti-TROP-2 cathepsin S-selective ADCs (313 and 314) as well as P1 _D_-amino acid non-cleavable controls (313D and 314D) in TROP-2^+^ MDA-MB-231 cells. The y-axis represents cell viability (%), and the x-axis shows ADC concentration on a log10 scale, ranging from 0.1 nM to 10 µM.

### Single-cell analysis of cathepsin S expression in breast tumors by mass cytometry (CyTOF)

Multiple studies have demonstrated elevated cathepsin S expression and activity in aggressive breast cancer subtypes, particularly in triple-negative breast cancer and HER-2-positive tumors (5, 7, 8). Immunohistochemical analyses of patient cohorts have shown that this protease is expressed in the majority of breast tumors, with the highest levels observed in high-grade, ER-negative, HER-2-positive, and TNBC cases (8). While HER-2 has been successfully exploited as a target antigen for several clinically approved ADCs, a substantial subset of ER-negative breast cancers, including many TNBCs, do not overexpress HER-2 and are therefore ineligible for HER-2-directed therapies. In these tumors, alternative cell-surface antigens are required to enable ADC-based strategies. TROP-2, a transmembrane glycoprotein overexpressed in a broad spectrum of epithelial malignancies, has emerged as a promising target for ADC development in breast cancer, particularly in HER-2-low or HER-2-negative disease (52, 53). The clinical activity of TROP-2-directed ADCs in heavily pretreated breast cancer underscores its translational relevance as a complementary or alternative target to HER-2. From a biomarker perspective, integrating information on TROP-2 expression with protease profiles such as cathepsin S may facilitate rational selection of both the target antigen and the protease-cleavable linker. Such an approach could extend the benefits of protease-activated ADCs to patient subsets that lack HER-2 but exhibit robust TROP-2 expression. Importantly, cathepsin S is secreted not only by malignant epithelial cells but also by tumor-infiltrating immune cells, especially macrophages and dendritic cells (54). High cathepsin S expression in stromal (infiltrating) cells has been associated with poor clinical outcomes, consistent with its role in tumor-promoting macrophage activity and extracellular matrix remodeling (8). In contrast, high cathepsin S expression within tumor epithelial cells, particularly in TNBC, correlates with improved prognosis and increased polarization of macrophages toward the M1 phenotype, suggesting a context-dependent, potentially protective function in this compartment (7). This dual role highlights the critical importance of spatial and cellular localization when evaluating cathepsin S as a biomarker. Furthermore, epithelial cathepsin S expression in TNBC has been linked to tumor subtypes with deficiencies in DNA damage repair pathways, indicating potential predictive value for sensitivity to DNA-damaging chemotherapies and reinforcing the need for further exploration of this protease in ER-negative breast cancer (7). To investigate cathepsin expression at single-cell resolution, we applied mass cytometry (CyTOF) to primary breast tumor samples. Mass cytometry is a high-dimensional proteomic technology that enables simultaneous quantification of dozens of protein markers in individual cells, offering a powerful platform for characterizing tumor composition and microenvironmental heterogeneity (55, 56). Although CyTOF is widely used to study immune infiltration, cell state dynamics, and therapeutic responses (57), its application to profiling protease expression, particularly in the context of protease-activated antibody-drug conjugates, remains underexplored. Given the growing interest in tumor-selective, protease-cleavable linkers for targeted therapies, mass cytometry provides a unique opportunity to assess the cellular and spatial distribution of activating enzymes such as cathepsin S directly in patient-derived tissues. In this study, we leveraged CyTOF to map cathepsin S expression across diverse cell populations within breast tumors, generating translational insights into its potential as a selective trigger for next-generation protease-activated therapeutics.

We performed mass cytometry (CyTOF) analysis on freshly resected tumor samples obtained from four patients diagnosed with grade 2 breast cancer. These patients displayed varying expression levels of HER-2, progesterone receptor (PR), and estrogen receptor (ER) (**Figure 6A**). The tumor tissues were processed into single-cell suspensions and stained with a previously developed panel of metal-conjugated antibodies (**Figure 6B**)(27). This panel enabled high-dimensional profiling of the tumor architecture, allowing for the identification of major cell types including epithelial and endothelial cells, fibroblasts, and various immune populations such as T cells, B cells, NK cells, and macrophages. In addition to lineage markers, the panel also included antibodies against key breast cancer biomarkers (HER-2, ER, PR) and cathepsin S. Dimensionality reduction using viSNE revealed significant inter-patient heterogeneity in tumor composition (**Figure 6C**). Two of the samples were dominated by epithelial and endothelial cells, while the other two showed substantial fractions of fibroblasts and infiltrating immune cells. HER-2 expression was largely confined to epithelial and endothelial populations, though its intensity and distribution varied among patients (**Figure 6D**). Notably, HER-2 expression only partially overlapped with ER and PR, and although some tumors expressed all three markers, their localization was distinct (**Figure S4**). However, due to the limited number of analyzed samples, these trends remain preliminary. Another biomarker currently under clinical evaluation for antibody-drug conjugate development is TROP-2 (52, 58). TROP-2 expression was detected in three of four patients, with the strongest epithelial expression observed in sample MASP-54 (**Figure 6E**). The other two samples also showed epithelial expression, but at lower abundance. Analysis of cathepsin S expression showed that in HER-2-positive tumors, this protease was co-expressed with HER-2 at the single-cell level and primarily localized to epithelial and endothelial compartments (**Figure 6F**). This spatial correlation suggests that cathepsin S may serve as a suitable proteolytic trigger for HER-2-targeting antibody-drug conjugates, enabling dual-selective therapeutic activation. Interestingly, in the triple-negative breast cancer sample (patient MASP55), cathepsin S expression was not detected in cancer epithelial cells but was abundant in infiltrating immune cells, particularly lymphocytes. This observation aligns with previous studies highlighting immune cell-derived cathepsin S as a significant contributor to the tumor protease landscape (reviewed in (5, 59, 60)). Moreover, in sample MASP-54, TROP-2, but not HER-2, correlated strongly with cathepsin S, suggesting that optimal protease-biomarker pairing for efficient ADC activation may vary among patients, and providing a tissue-level analogue of the TROP-2-cathepsin S axis functionally validated by our sacituzumab-based ADCs in MDA-MB-231 cells. As expected, cathepsin B was highly abundant in epithelial cells, and to a lesser extent in immune cells, in patients MASP54, MASP55, and MASP57 (**Figure 6G**). However, to our surprise, cathepsin B was essentially absent in MASP52. Interestingly, in MASP52 cathepsin S was readily detected and co-localized with HER-2 in epithelial clusters, whereas cathepsin B did not. This profile indicates HER-2-cathepsin S as the optimal trigger pair for linker design in this tumor. Although epithelial dominance of cathepsin S over cathepsin B appears to be rare in breast cancer, MASP52 provides a clear example where protease selection should follow the biomarker rather than default practice. These data support biomarker-protease pairing in personalized diagnostics and suggest that patients with low tumoral cathepsin B may experience suboptimal responses to cathepsin B-cleavable linkers. Our findings have important implications for the design of CTSS-activated therapeutics. In TNBC, where tumor cells may lack suitable surface markers for targeting, extracellular activation by infiltrating immune-cell-derived proteases like cathepsin S may offer a viable alternative. Our demonstration that HER-2-negative, TROP-2-positive MDA-MB-231 cells can be efficiently killed by sacituzumab-based, cathepsin S-cleavable ADCs illustrates how such protease-guided strategies can be combined with alternative antigens when they are present, while still allowing for non-internalizing, microenvironment-driven activation in antigen-poor tumors. This strategy could enable the development of non-internalizing ADCs or peptide-based prodrugs that are activated in the tumor microenvironment, generating cytotoxic effects via a bystander mechanism (61). Given the known heterogeneity of tumor marker expression and the frequent absence of internalizing targets in TNBC, extracellular protease activation may provide a more robust and broadly effective approach for treating these aggressive cancers.

**Figure 6.**
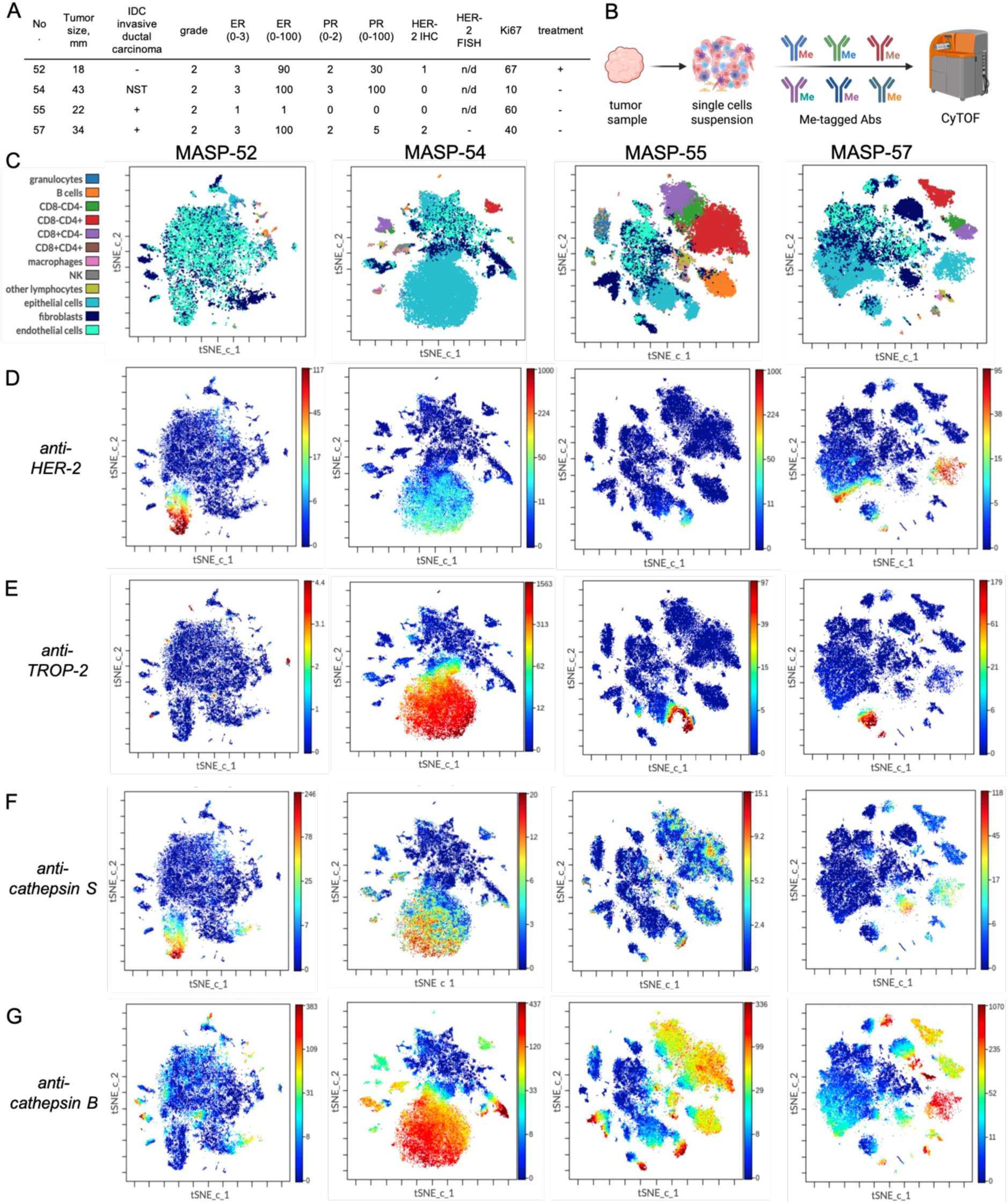
Single-cell mass cytometry profiling of the breast tumor microenvironment. **(A)** Freshly resected breast tumor tissues were enzymatically dissociated into single-cell suspensions and analyzed by mass cytometry (CyTOF) to generate a high-dimensional cellular landscape of the tumor microenvironment. (**B**) Clinical characteristics of breast cancer patients included in the CyTOF analysis. The table summarizes key pathological features, including tumor size, histological grade, estrogen receptor (ER) and progesterone receptor (PR) status, HER-2 expression, and the proliferation marker Ki-67. **(C)** viSNE analysis revealed extensive intratumoral heterogeneity across patient samples. While epithelial cells, endothelial cells, and fibroblasts were the dominant populations in all tumors, variable levels of immune cell infiltration, including T cells, B cells, macrophages, and NK cells, were detected in individual patients. **(D,E,F,G)** viSNE-based visualization of HER-2 (**D**), TROP-2 (**E**), cathepsin S (**F**), and cathepsin B **(G)** expression across diverse tumor cell populations. Expression levels are shown as heat maps, with red indicating high expression and blue indicating low or no expression.

## Conclusions

In this study, we developed a chemical toolkit towards cathepsin S, including highly selective substrates, irreversible inhibitors, and activity-based probes. We also integrated a cathepsin S-selective peptide sequence into a cathepsin S-susceptible linker for newly engineered ADCs, which demonstrated promising cytotoxic activity in a TNBC model. Using the HyCoSuL approach with a broad panel of unnatural amino acids, we systematically profiled cathepsin S substrate preferences and engineered optimized peptide sequences with superior selectivity and catalytic efficiency. These substrates enabled the identification of key structural motifs that guided the rational design of tailored inhibitors and ABPs. The resulting AOMK-based inhibitors demonstrated potent, time-dependent inhibition of cathepsin S while sparing related proteases, cathepsins L, B and V. Their specificity made them ideal scaffolds for ABPs, which we functionalized with fluorophores isotopes for use in fluorescence imaging and mass cytometry, respectively. These probes provided valuable tools for assessing cathepsin S activity at the biochemical and cellular levels and will be instrumental for future studies of tumor protease landscapes. Together, these optimized substrates and ABP-based activity studies established both the selective peptide motifs and the biological context required to translate our findings into cathepsin S-cleavable therapeutic designs. Therefore, building on these optimized sequences, we developed the first cathepsin S-cleavable MMAE prodrugs and corresponding antibody-drug conjugates. These constructs showed enzyme-dependent activation, strong cytotoxicity in cathepsin S-positive MDA-MB-231 cells, and minimal off-target effects in cathepsin S-negative cell lines. In ADC formats, the selective linker enabled payload release specifically in HER-2^+^ tumors expressing cathepsin S and, when combined with sacituzumab, in HER-2-negative/TROP-2-positive TNBC cells, validating the therapeutic utility of this protease as a tumor-activating trigger across molecularly distinct breast cancer subtypes. Importantly, the kinetic characterization of our lead substrates underscores how distinct enzymatic parameters can influence antibody-drug conjugate linker performance. Notably, **OG-197** (Phe(F_5_)-Cit-Lys(2ClZ)-Glu(Me)) and **OG-209** (Met(O_2_)-Cit-NptGly-Glu(Me)) substrates achieved similar catalytic efficiencies (k_cat_/KM of 356,000 and 564,000 M^−1^s^−1^, respectively) for cathepsin S, yet they did so via different strategies. **OG-197** relies on an exceptionally high affinity (K_M_ of 0.74 µM) coupled with a modest turnover rate (k_cat_ 0.26 s^−1^). In contrast, **OG-209** sacrifices binding strength (K_M_ of 39,2 µM) for a vastly higher turnover (k_cat_ of 22,1 s^−1^). Importantly, both peptide sequences remain highly selective for cathepsin S over other cathepsins, confirming that our HyCoSuL-derived motifs intrinsically favor cathepsin S. This selectivity is reflected in cell-free assays; however, within cellular environments we observed that activity-based probes (ABPs) derived from these sequences (e.g., **OG-233**, **OG-234**) can exhibit minor off-target labeling of cathepsin B, likely due to the overwhelming abundance of cathepsin B in certain cancer cells. Such findings highlight that probe labeling patterns in cells are influenced by protease expression levels, whereas the substrate kinetics demonstrate true enzyme preference. From a prodrug activation standpoint, a low K_M_ may be especially advantageous under the nanomolar substrate concentrations encountered after ADC internalization. In these conditions, **313**’s tighter binding ensures that cathepsin S actively engages the linker even at low intracellular drug levels, favoring efficient payload release. By contrast, a high K_M_ linker like **314** might require higher local concentrations to achieve comparable enzyme occupancy, potentially delaying drug liberation. Thus, for cathepsin S-activated ADCs, optimizing substrate affinity (K_M_) can be just as critical as maximizing k_cat_, ensuring that even a few intracellular prodrug molecules are rapidly bound and cleaved by the target protease in the competitive milieu of the lysosome. Moreover, cathepsin S possesses unique properties that make it particularly suitable for targeted drug activation. Unlike most cathepsins, cathepsin S is active at neutral pH and functions not only intracellularly but also in the extracellular tumor microenvironment. Therefore, by exploiting this unique stability and activity at neutral pH we unlock the potential for extracellular drug release mechanisms. Cathepsin S is known to be secreted by tumor-associated cells and remains active in the slightly acidic to neutral tumor microenvironment. Thus, in addition to conventional internalization-dependent payload release, a cathepsin S-cleavable ADC could even be activated in the extracellular milieu. This opens the door to “non-internalizing” ADC or prodrug strategies, whereby a tumor-targeted antibody-drug conjugate need not enter every cell to be effective -the protease-rich microenvironment can trigger drug release outside cells, enabling a potent bystander effect that kills adjacent tumor cells. Together, these attributes emphasize how a cathepsin S-selective linker strategy can broaden the therapeutic window: combining high tumor specificity with versatile activation modes to address the heterogeneity of tumor biology.

From a diagnostic perspective, cathepsin S is upregulated in aggressive tumor subtypes such as triple-negative breast cancer and shows limited expression in most normal tissues, supporting its role as a selective tumor activator. To assess the relevance of cathepsin S expression in clinical samples, we applied mass cytometry to analyze single-cell protein profiles in breast tumors. This profiling revealed striking inter-patient variability in protease expression, reinforcing the need for personalized ADC linker strategies. In three of the four patient samples analyzed, cathepsin B was the dominant cysteine protease in tumor cells (with cathepsin S either lower in abundance or largely restricted to infiltrating immune cells). However, one tumor (patient MASP52) exhibited the opposite pattern -high cathepsin S expression coupled with negligible cathepsin B. This outlier finding is clinically significant: a “one-size-fits-all” ADC employing a cathepsin B-cleavable linker (e.g. the prevalent Val-Cit motif, which is broadly active but non-specific) might underperform in such a patient, as the intended activating enzyme is scarce. Conversely, a cathepsin S-cleavable ADC would be ideally suited to exploit the available proteolytic machinery in MASP52’s tumor. More generally, these data suggest that matching the ADC linker to the tumor’s protease profile could maximize therapeutic efficacy. In HER-2-positive tumors, we observed cathepsin S co-expressed with HER-2 in the same cell populations, implying that a HER-2-targeted ADC with a cathepsin S-sensitive linker could achieve dual selectivity: first by antibody-antigen binding, and second by protease-specific payload release within those HER-2^+^ cancer cells. In parallel, our CyTOF data and functional studies with sacituzumab-based ADCs demonstrate that TROP-2 provides a complementary antigenic handle in HER-2-low or HER-2-negative settings, particularly when co-expressed with cathepsin S at the single-cell level. In patient subsets where cathepsin B is abundant, traditional cathepsin B-cleavable linkers may suffice, but in subsets exemplified by MASP52 (high cathepsin S/low cathepsin B), a cathepsin S-triggered ADC linker may significantly outperform the conventional design. These results underscore the importance of protease profiling as a companion diagnostic. By tailoring the cleavage mechanism of an ADC to the proteolytic landscape of a patient’s tumor, one can enhance selective drug activation and potentially improve clinical outcomes in heterogeneous diseases like breast cancer.

Collectively, our results highlight the power of unnatural amino acid-based chemistry to engineer highly selective protease-targeting reagents. The cathepsin S-specific substrates, inhibitors, and ABPs developed here lay the foundation for broad applications in protease biology and diagnostics. In parallel, our selective prodrugs and ADCs demonstrate the feasibility of using cathepsin S as a tumor-restricted activator of cytotoxic therapies. Finally, mass cytometry provides a critical diagnostic platform to match these targeted therapeutics with appropriate patient profiles, advancing the concept of protease-guided precision oncology.

## Acknowledgements

This project was supported by the National Science Centre in Poland; grant SONATA UMO-2018/31/D/NZ5/02406 to MP and OPUS-LAP UMO-2020/39/I/NZ5/03104 to MP, and grants P1-0140 and N1-0229 to BT.

## Authors contribution

M.Ł. conceptualized the *in vitro* ADC studies, synthesized antibody-drug conjugates, performed the *in vitro* analysis of cathepsin S peptide prodrugs and ADCs in breast cancer cells, performed labeling of active cathepsins using activity-based probes (ABPs) and wrote the draft of the manuscript. O.G., synthesized cathepsin substrates, ABPs and peptide prodrugs, performed LC-MS analyses, synthesized antibody-drug conjugates, and carried out *in vitro* cytotoxicity assays in breast cancer cell lines. N.Ć.-P. performed all mass cytometry (CyTOF) experiments on breast tumor samples. J.N. and V.P. synthesized cathepsin S inhibitors and performed their kinetic analysis. M.M. performed the synthesis of peptide prodrugs and antibody-drug conjugates. P.K. collected and processed breast tumor samples. B.D.-K. collected and processed breast tumor samples. B.T. provided recombinant cathepsins and contributed to experimental design, data analysis, and interpretation for protease-related assays. M.D. provided P1-Gln and P1-Arg HyCoSuL library and contributed to data analysis. R.M. collected and processed breast tumor samples, supervised the experimental design, and contributed to data analysis and interpretation. M.P. conceptualized and supervised the study, designed the experiments, performed screening of cathepsin S substrate specificity, led peptide prodrug development, provided funding and resources, and wrote the draft of the manuscript. All authors critically reviewed, revised, and approved the final manuscript.

