## Supplemental File for "Engineering cathepsin S selective chemical probes and antibody-drug conjugates through substrate profiling with unnatural amino acids"

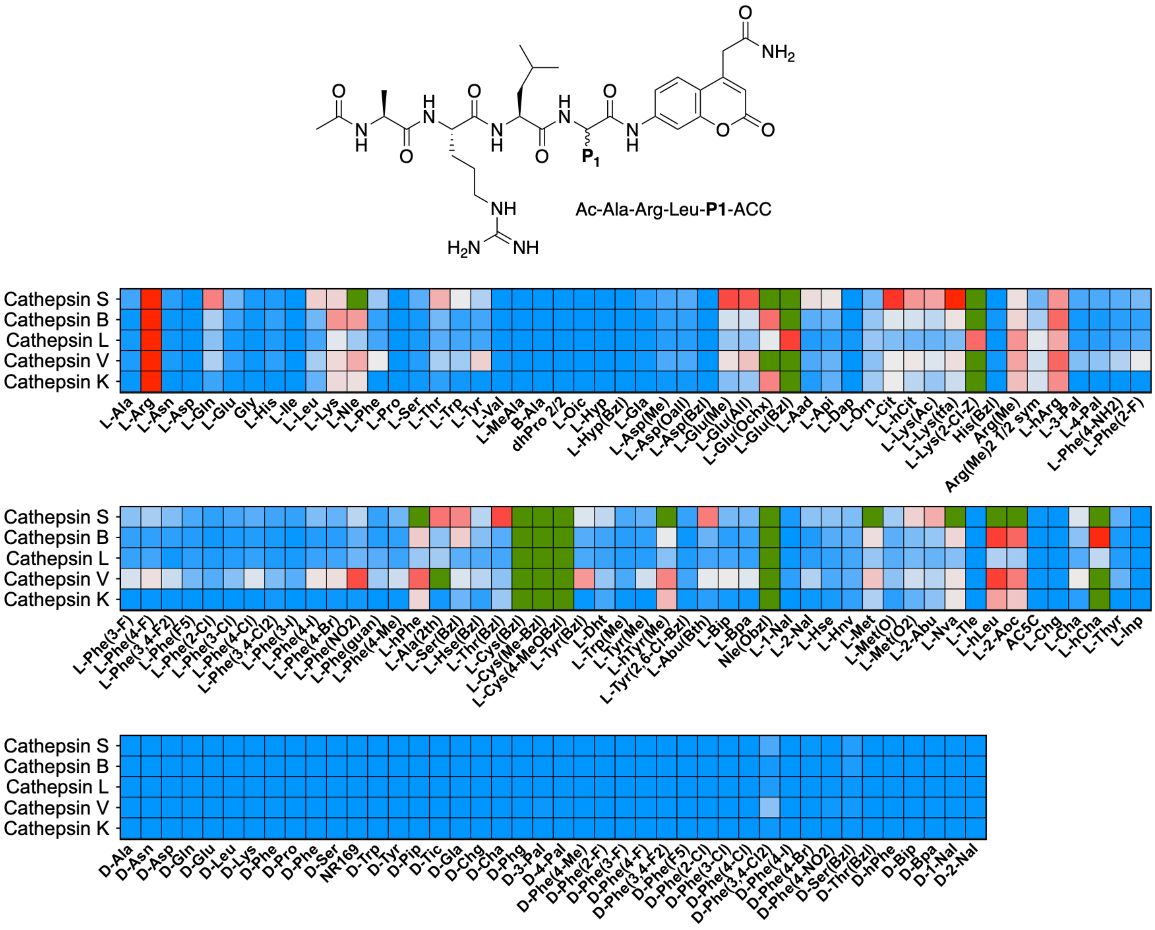


**Figure S1 Substrate specificity of cysteine cathepsins at the P1 position.** The P1 substrate preferences of human cysteine cathepsins were determined using a fluorogenic substrate library based on the Ac-Ala-Arg-Leu-P1-ACC scaffold, comprising 19 natural and over 100 unnatural amino acids. Specificity profiling was performed in triplicate. Substrate hydrolysis rates (RFU/s) are presented as average values, with standard deviations (SD) below 10% for each substrate. To facilitate comparison, data were normalized to the activity against arginine (set to 100%), and a red-to-blue heatmap was used to visualize relative preferences. Unnatural amino acids that were better recognized than arginine (>100%) are highlighted in green. P1 specificity data for cathepsins B, L, V, and K were taken from our previously published studies.

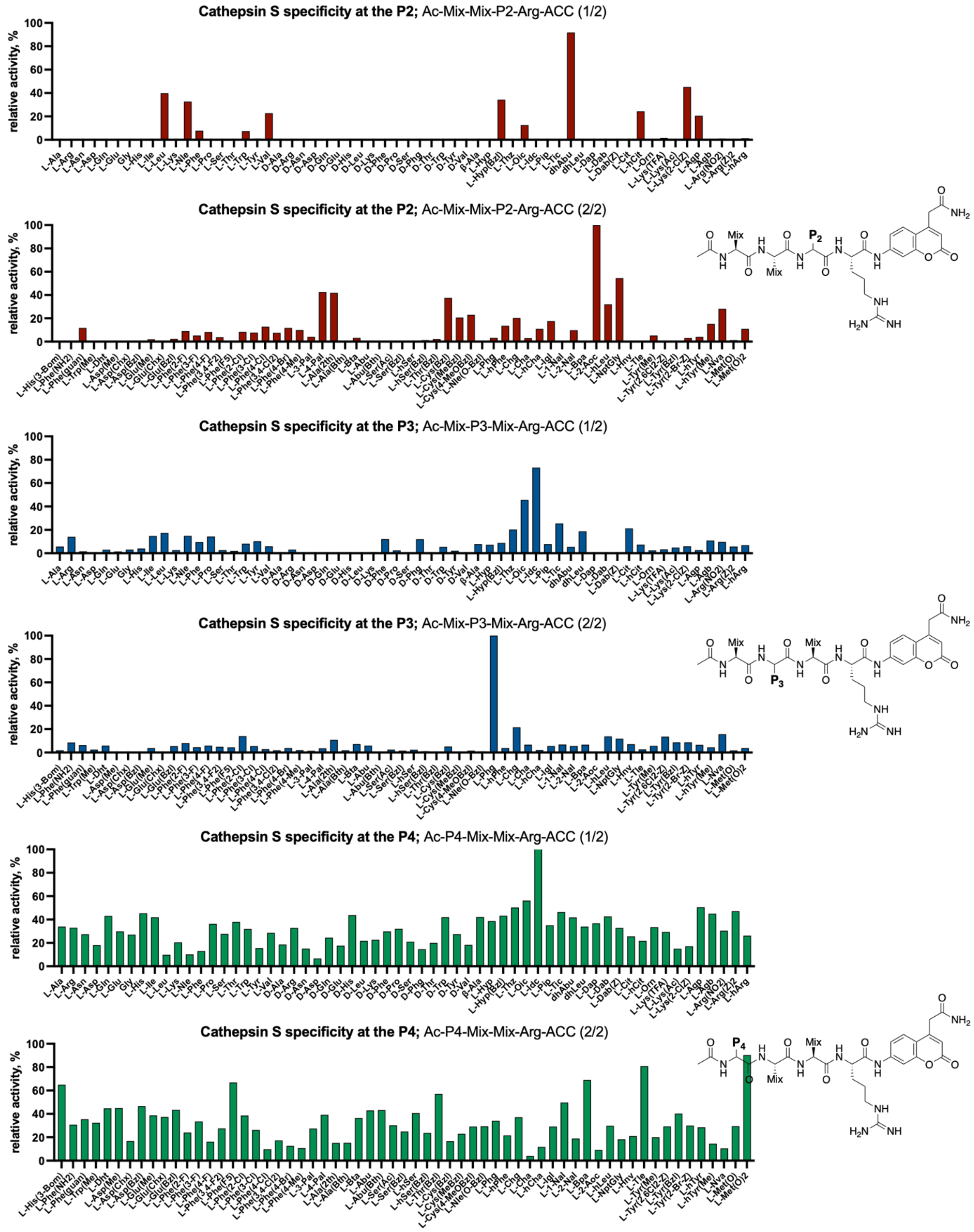


**Figure S2 Substrate specificity of cathepsin S at the P4-P2 positions (P1-Arg HyCoSuL).** The x-axis shows the abbreviated names of natural and unnatural amino acids, while the y-axis represents the relative enzymatic activity of cathepsin S for each substrate. The most efficiently cleaved amino acid at each position was set to 100%, and the activities of other amino acids were normalized accordingly. The structures of the P4, P3, and P2 sublibraries are presented on the right side.

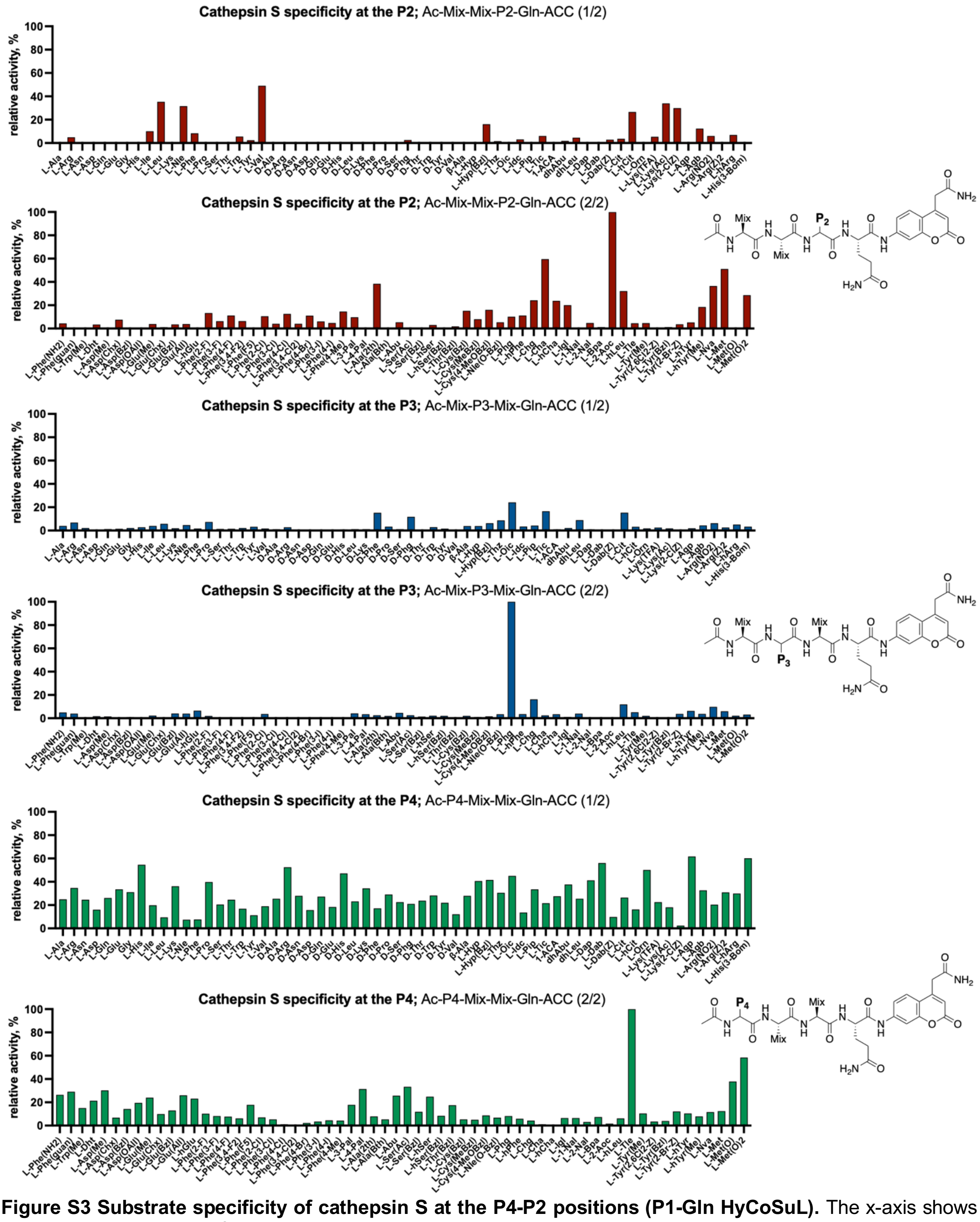


**Figure S3 Substrate specificity of cathepsin S at the P4-P2 positions (P1-Gln HyCoSuL).** The x-axis shows the abbreviated names of natural and unnatural amino acids, while the y-axis represents the relative enzymatic activity of cathepsin S for each substrate. The most efficiently cleaved amino acid at each position was set to 100%, and the activities of other amino acids were normalized accordingly. The structures of the P4, P3, and P2 sublibraries are presented on the right side.

**Table S1 Chemical structures of first-generation substrates for cathepsin S.** All substrates follow the general structure Ac-P4-P3-P2-Arg-ACC. The table lists the sequences and corresponding chemical structures designed to explore substrate preferences at the P4-P2 positions, with a fixed Arg residue at P1.

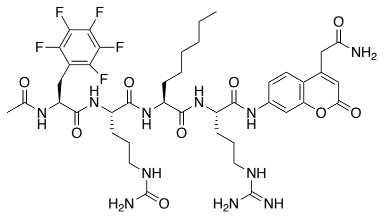

**OG-64**
Ac-Phe($F_5$)-Cit-2Aoc-Arg-ACC

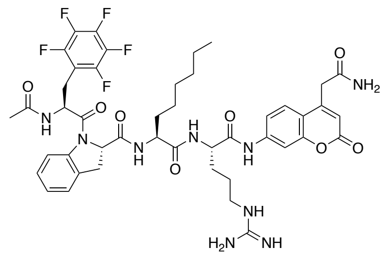

**OG-65**
Ac-Phe($F_5$)-Idc-2Aoc-Arg-ACC

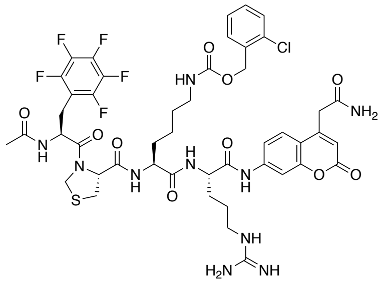

**OG-66**
Ac-Phe($F_5$)-Thz-Lys(2ClZ)-Arg-ACC

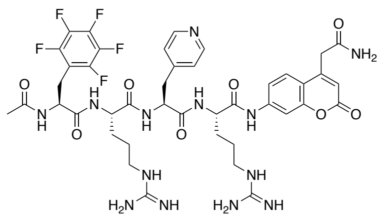

**OG-67**
Ac-Phe($F_5$)-Cit-3Pal-Arg-ACC

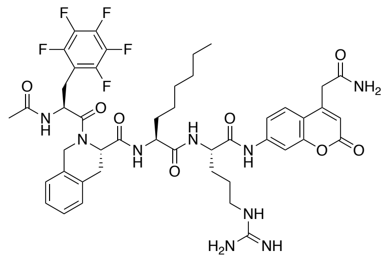

**OG-68**
Ac-Phe($F_5$)-Tic-2Aoc-Arg-ACC

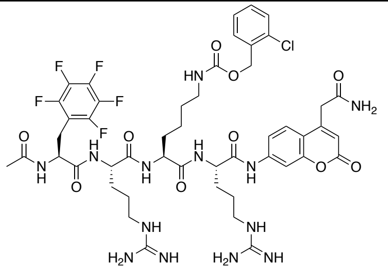

**OG-69**
Ac-Phe($F_5$)-Arg-Lys(2ClZ)-Arg-ACC

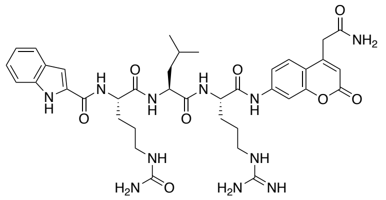

**OG-70**
HN-Idc-Cit-Leu-Arg-ACC

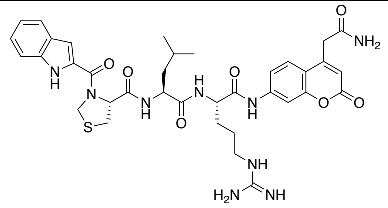

**OG-71**
HN-Idc-Thz-Leu-Arg-ACC

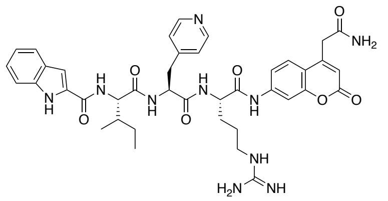

**OG-72**
HN-Idc-Ile-3Pal-Arg-ACC

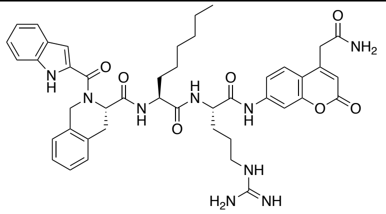

**OG-73**
HN-Idc-Tic-2Aoc-Arg-ACC

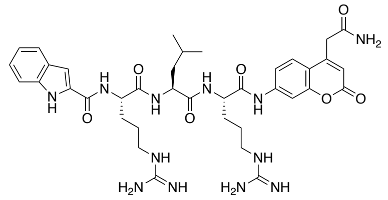

**OG-74**
HN-Idc-Arg-Leu-Arg-ACC

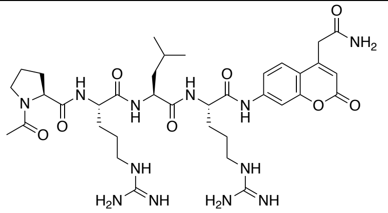

**OG-75**
Ac-Pro-Arg-Leu-Arg-ACC

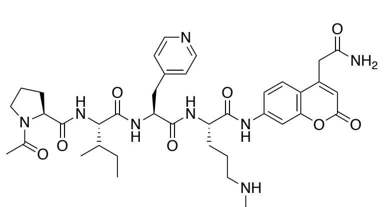

**OG-76**
Ac-Pro-Ile-3Pal-Arg-ACC

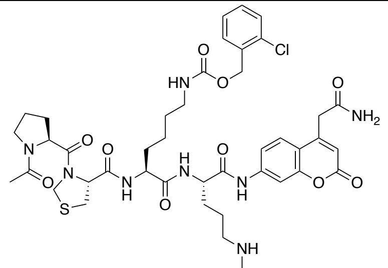

**OG-77**
Ac-Pro-Thz-Lys(2ClZ)-Arg-ACC

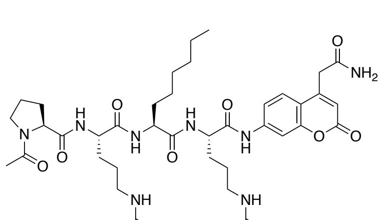

**OG-79**
Ac-Pro-Arg-2Aoc-Arg-ACC

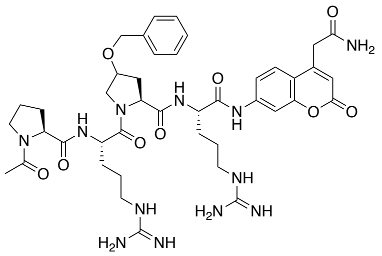

**OG-80**
Ac-Pro-Arg-Hyp(Bzl)-Arg-ACC

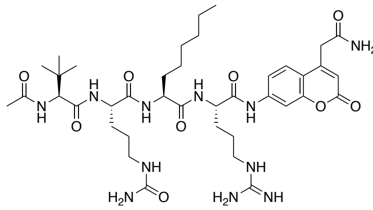

**OG-81**
Ac-Tle-Cit-2Aoc-Arg-ACC

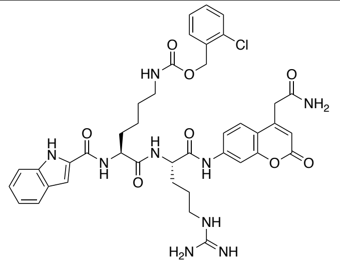

**OG-82**
Ac-Tle-Idc-Lys(2ClZ)-Arg-ACC

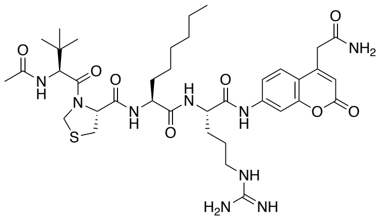

**OG-83**
Ac-Tle-Thz-2Aoc-Arg-ACC

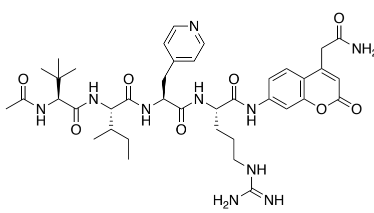

**OG-84**
Ac-Tle-Ile-3Pal-Arg-ACC

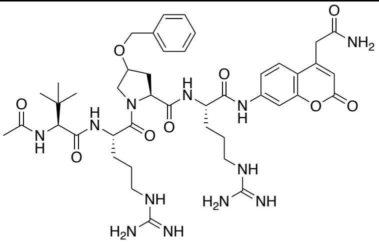

**OG-85**
Ac-Tle-Arg-Hyp(Bzl)-Arg-ACC

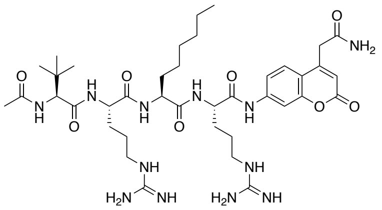

**OG-86**
Ac-Tle-Arg-2Aoc-Arg-ACC

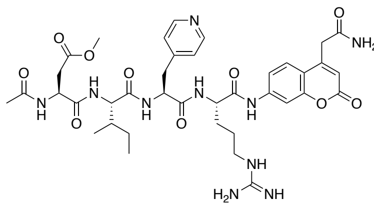

**OG-87**
Ac-Asp(Me)-Ile-3Pal-Arg-ACC

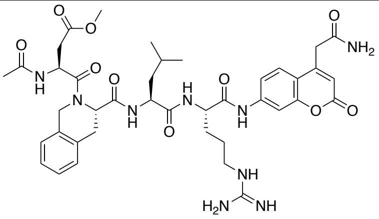

**OG-88**
Ac-Asp(Me)-Tic-Leu-Arg-ACC

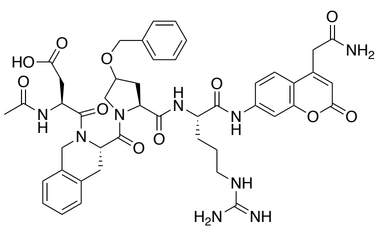

**OG-89**
Ac-Asp(Bzl)-Tic-Hyp(Bzl)-Arg-ACC

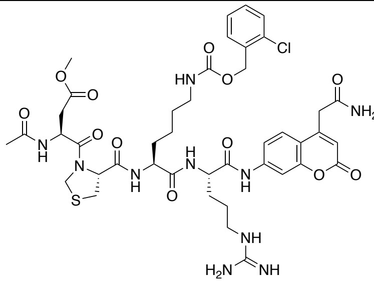

**OG-90**
Ac-Asp(Bzl)-Thz-Lys(2ClZ)-Arg-ACC

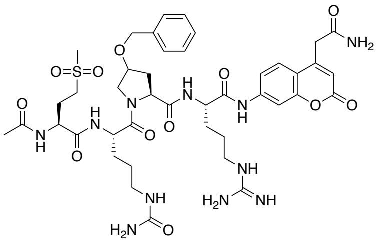

**OG-91**
Ac-Met($O_2$)-Cit-Hyp(Bzl)-Arg-ACC

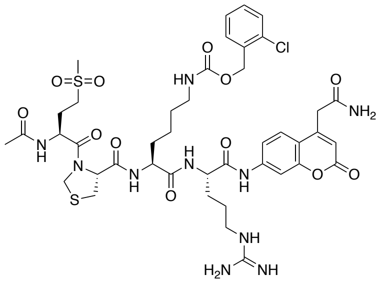

**OG-93**
Ac-Met($O_2$)-Thz-Lys(2ClZ)-Arg-ACC

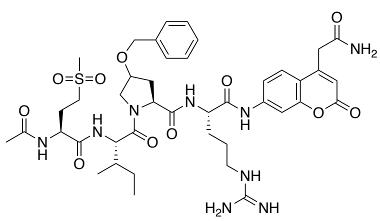

**OG-94**
Ac-Met($O_2$)-Ile-Hyp(Bzl)-Arg-ACC

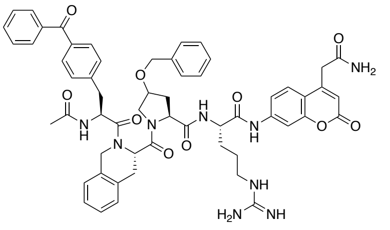

**OG-95**
Ac-Bpa-Tic-Hyp(Bzl)-Arg-ACC

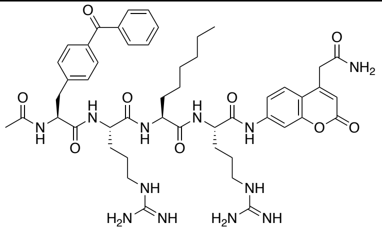

**OG-96**
Ac-Bpa-Arg-2Aoc-Arg-ACC

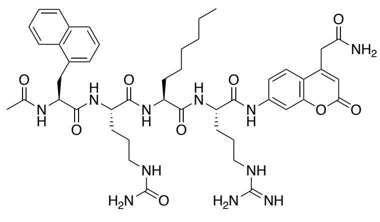

**OG-115**
Ac-1Nal-Cit-2Aoc-Arg-ACC

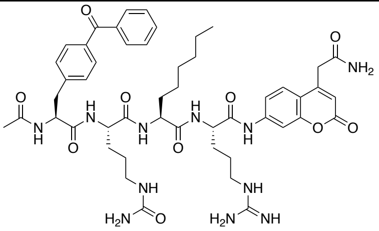

**OG-116**
Ac-Bpa-Cit-2Aoc-Arg-ACC

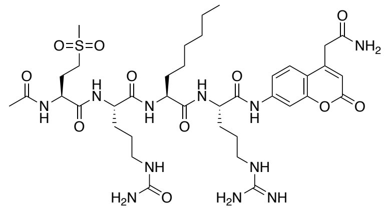

**OG-117**
Ac-Met($O_2$)-Cit-2Aoc-Arg-ACC

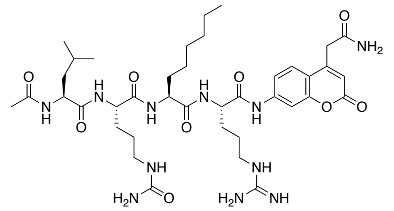

**OG-118**
Ac-Leu-Cit-2Aoc-Arg-ACC

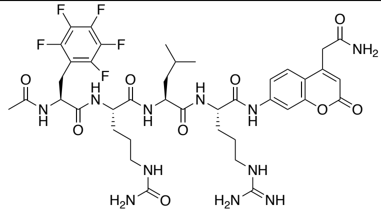

**OG-119**
Ac-Phe($F_5$)-Cit-Leu-Arg-ACC

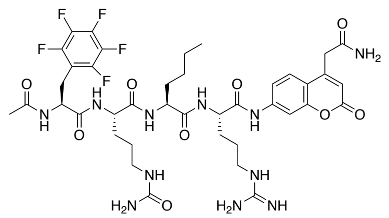

**OG-120**
Ac-Phe($F_5$)-Cit-Nle-Arg-ACC

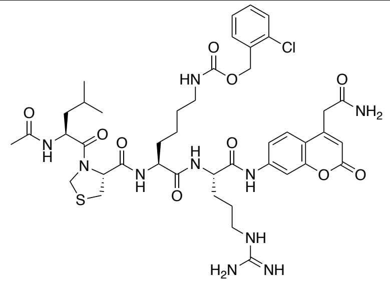

**OG-121**
Ac-Leu-Thz-Lys(2ClZ)-Arg-ACC

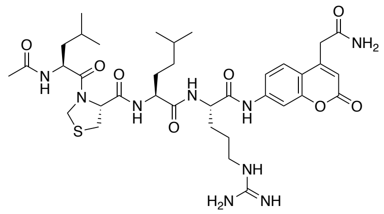

**OG-122**
Ac-Leu-Thz-hLeu-Arg-ACC

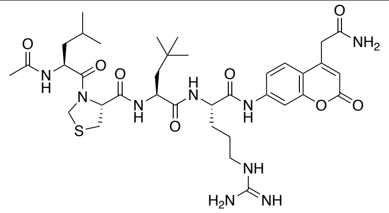

**OG-123**
Ac-Leu-Thz-NptGly-Arg-ACC

**Table S2 Chemical structures of second-generation substrates for cathepsin S.** All substrates follow the general structure Ac-Phe($F_5$)-Cit-2Aoc-P1-ACC. This library was designed to optimize the P1 position using the fixed P4-P2 motif derived from the most selective first-generation substrate.

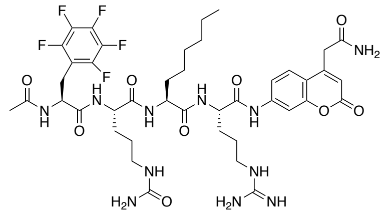

**OG-64**
Ac-Phe($F_5$)-Cit-2Aoc-Arg-ACC

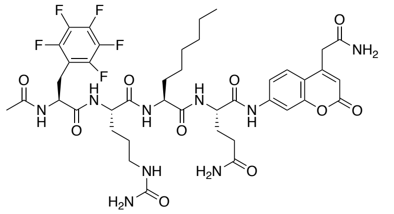

**OG-133**
Ac-Phe($F_5$)-Cit-2Aoc-Gln-ACC

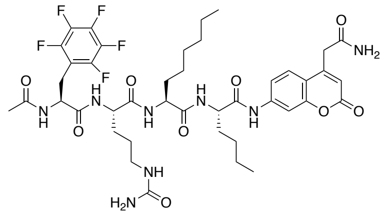

**OG-134**
Ac-Phe($F_5$)-Cit-2Aoc-Nle-ACC

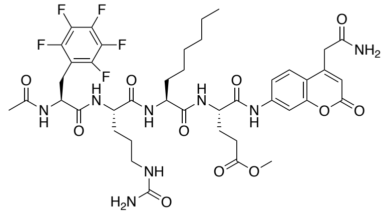

**OG-135**
Ac-Phe($F_5$)-Cit-2Aoc-Glu(Me)-ACC

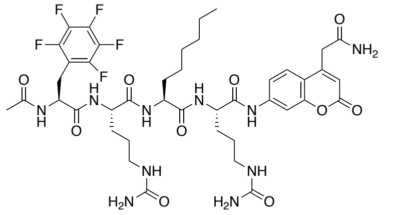

**OG-136**
Ac-Phe($F_5$)-Cit-2Aoc-Cit-ACC

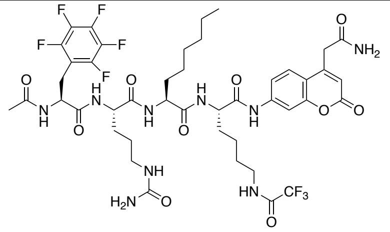

**OG-137**
Ac-Phe($F_5$)-Cit-2Aoc-Lys(TFA)-ACC

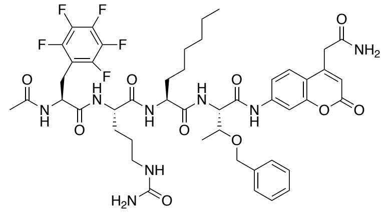

**OG-138**

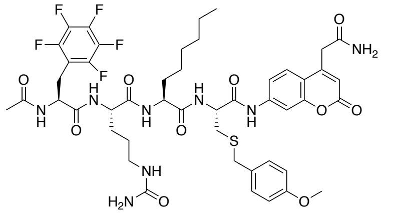

**OG-139**

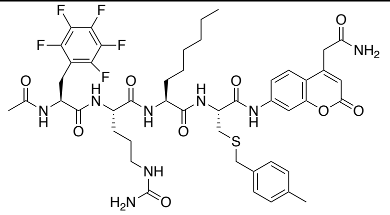

**OG-140**

| | | |
|---|---|---|
| Ac-Phe($F_5$)-Cit-2Aoc-Thr(Bzl)-ACC | Ac-Phe($F_5$)-Cit-2Aoc-Cys(MeOBzl)-ACC | Ac-Phe($F_5$)-Cit-2Aoc-Cys(MeBzl)-ACC |
| 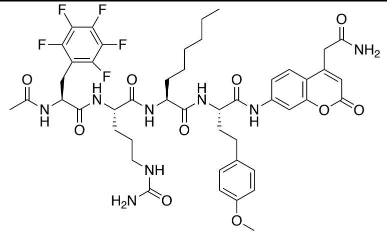 **OG-141** Ac-Phe($F_5$)-Cit-2Aoc-hTyr(Me)-ACC | 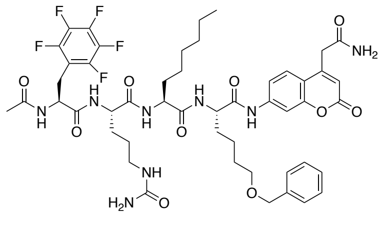 **OG-142** Ac-Phe($F_5$)-Cit-2Aoc-Nle(OBzl)-ACC | 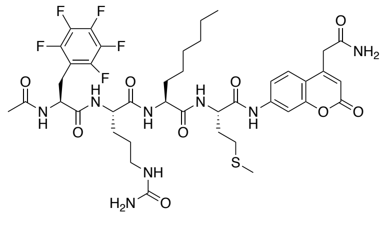 **OG-143** Ac-Phe($F_5$)-Cit-2Aoc-Met-ACC |
| 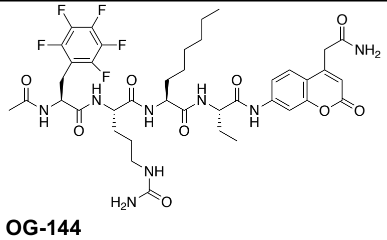 **OG-144** Ac-Phe($F_5$)-Cit-2Aoc-Abu-ACC | 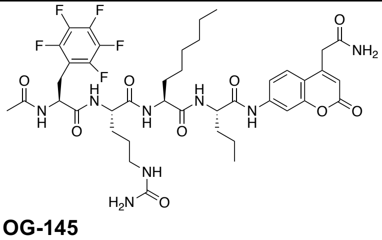 **OG-145** Ac-Phe($F_5$)-Cit-2Aoc-Nva-ACC | 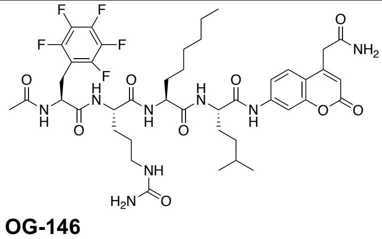 **OG-146** Ac-Phe($F_5$)-Cit-2Aoc-hLeu-ACC |
| 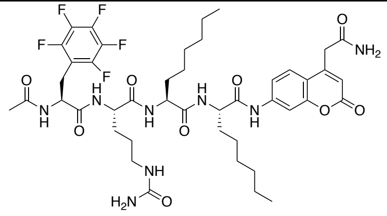 **OG-147** Ac-Phe($F_5$)-Cit-2Aoc-2Aoc-ACC | | |

**Table S3 Chemical structures of third-generation substrates for cathepsin S.** These substrates were developed by further refining the peptide sequence to enhance cathepsin S selectivity and activity. Variations in the P4-P1 positions were introduced based on previous substrate screening results, aiming to improve enzymatic cleavage efficiency and specificity.

| | | |
|---|---|---|
| 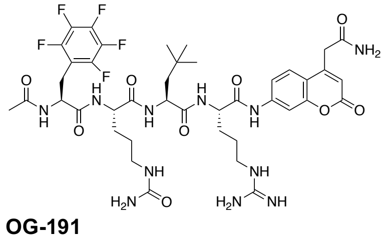 **OG-191** Ac-Phe($F_5$)-Cit-NptGly-Arg-ACC | 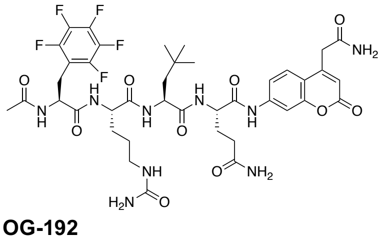 **OG-192** Ac-Phe($F_5$)-Cit-NptGly-Gln-ACC | 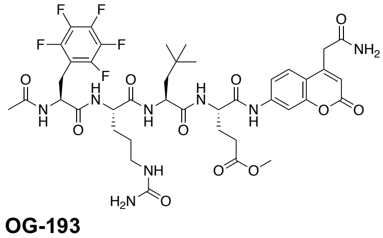 **OG-193** Ac-Phe($F_5$)-Cit-NptGly-Glu(Me)-ACC |
| 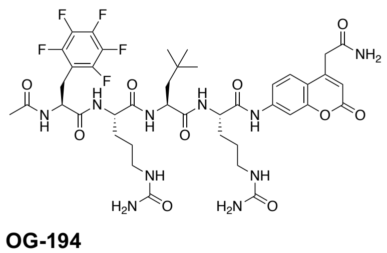 **OG-194** Ac-Phe($F_5$)-Cit-NptGly-Cit-ACC | 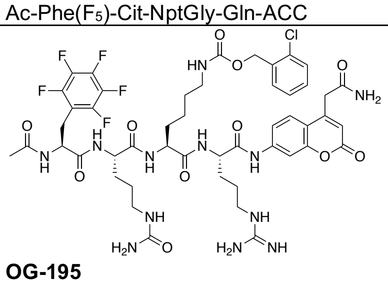 **OG-195** Ac-Phe($F_5$)-Cit-Lys(2ClZ)-Arg-ACC | 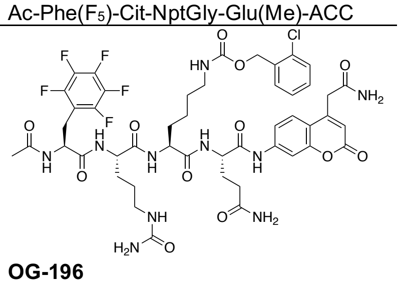 **OG-196** Ac-Phe($F_5$)-Cit-Lys(2ClZ)-Gln-ACC |

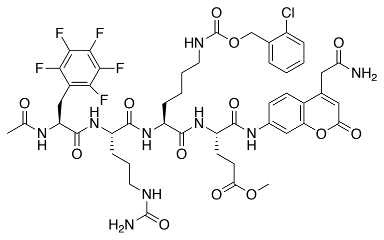

**OG-197**
Ac-Phe($F_5$)-Cit-Lys(2ClZ)-Glu(Me)-ACC

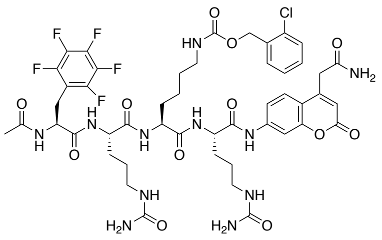

**OG-198**
Ac-Phe($F_5$)-Cit-Lys(2ClZ)-Cit-ACC

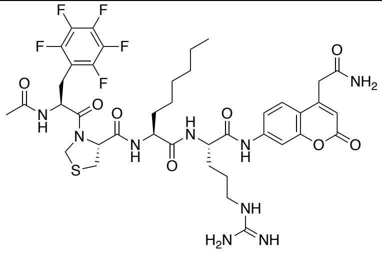

**OG-199**
Ac-Phe($F_5$)-Thz-2Aoc-Arg-ACC

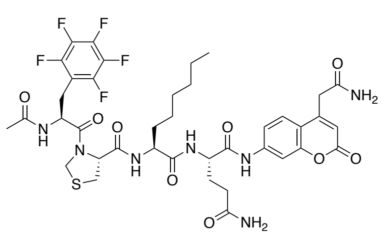

**OG-200**
Ac-Phe($F_5$)-Thz-2Aoc-Gln-ACC

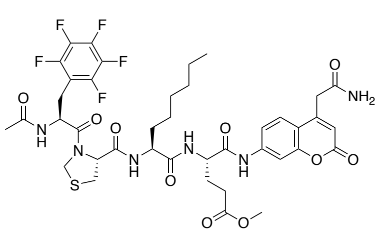

**OG-201**
Ac-Phe($F_5$)-Thz-2Aoc-Glu(Me)-ACC

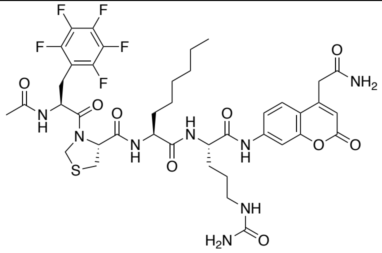

**OG-202**
Ac-Phe($F_5$)-Thz-2Aoc-Cit-ACC

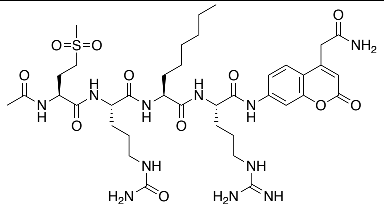

**OG-203**
Ac-Met($O_2$)-Cit-2Aoc-Arg-ACC

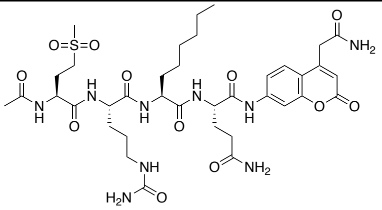

**OG-204**
Ac-Met($O_2$)-Cit-2Aoc-Gln-ACC

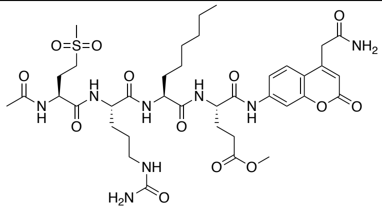

**OG-205**
Ac-Met($O_2$)-Cit-2Aoc-Glu(Me)-ACC

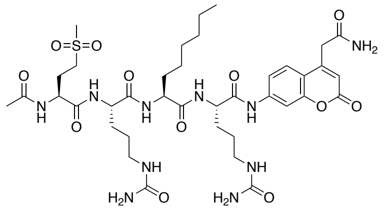

**OG-206**
Ac-Met($O_2$)-Cit-2Aoc-Cit-ACC

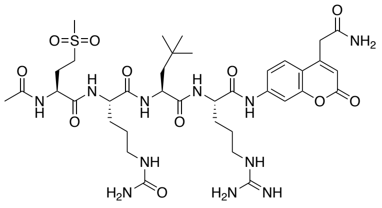

**OG-207**
Ac-Met($O_2$)-Cit-NptGly-Arg-ACC

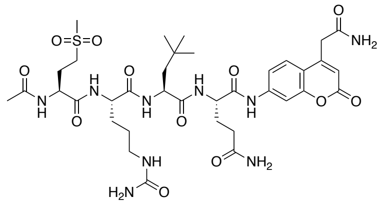

**OG-208**
Ac-Met($O_2$)-Cit-NptGly-Gln-ACC

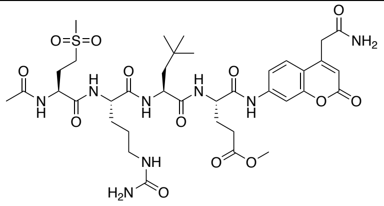

**OG-209**
Ac-Met($O_2$)-Cit-NptGly-Glu(Me)-ACC

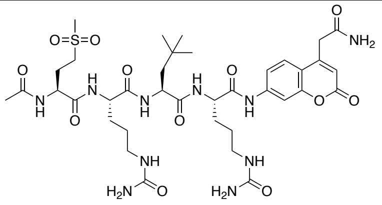

**OG-210**
Ac-Met($O_2$)-Cit-NptGly-Cit-ACC

**Table S4 Kinetic parameters ($k_{cat}/K_M$) of third-generation cathepsin S substrates.** Catalytic efficiency ($k_{cat}/K_M$) values were determined for each substrate to evaluate the impact of sequence modifications on cathepsin S cleavage. Measurements were performed under steady-state conditions, and the data reflect the enzyme's substrate preference and turnover efficiency.

| P4-P2 | P1 | Code | $k_{cat}/K_M$, $M^{-1}s^{-1}$ | | | |
|---|---|---|---|---|---|---|
| | | | Cathepsin S | Cathepsin B | Cathepsin L | Cathepsin V |
| Phe(F5)-Cit-2Aoc | Arg | **OG-64** | 574,000 | 144,000 | 13,800 | 25,400 |
| Phe(F5)-Cit-NptGly | Arg | **OG-191** | 382,000 | 32,200 | 5,000 | 92,800 |
| | Gln | **OG-192** | 119,000 | 6,400 | 6,100 | 28,100 |
| | Glu(Me) | **OG-193** | 133,000 | 7,900 | 7,800 | 22,700 |
| | Cit | **OG-194** | 115,000 | 11,200 | 6,700 | 38,600 |
| Phe(F5)-Cit-Lys(2ClZ) | Arg | **OG-195** | 364,000 | 73,200 | 10,200 | 14,200 |
| | Gln | **OG-196** | 688,000 | 45,400 | 5,200 | 10,400 |
| | Glu(Me) | **OG-197** | 356,000 | 5,500 | 2,100 | 7,700 |
| | Cit | **OG-198** | 438,000 | 18,700 | 4,400 | 17,900 |
| Phe(F5)-Thz-2Aoc | Arg | **OG-199** | 498,000 | 119,000 | 12,400 | 38,500 |
| | Gln | **OG-200** | 93,600 | 14,800 | 1,200 | 46,400 |
| | Glu(Me) | **OG-201** | 214,000 | 29,200 | 2,500 | 25,900 |
| | Cit | **OG-202** | 247,000 | 54,500 | 2,900 | 56,700 |
| Met(O2)-Cit-2Aoc | Arg | **OG-203** | 1,097,000 | 232,000 | 102,000 | 93,900 |
| | Gln | **OG-204** | 268,000 | 186,000 | 24,700 | 32,300 |
| | Glu(Me) | **OG-205** | 867,000 | 127,000 | 9,600 | 22,100 |
| | Cit | **OG-206** | 166,000 | 96,500 | 12,000 | 17,000 |
| Met(O2)-Cit-NptGly | Arg | **OG-207** | 529,000 | 21,900 | 50,700 | 83,800 |
| | Gln | **OG-208** | 91,400 | 3,700 | 8,900 | 36,800 |
| | Glu(Me) | **OG-209** | 564,000 | 5,100 | 8,300 | 33,100 |
| | Cit | **OG-210** | 219,000 | 7,500 | 9,800 | 37,200 |

**Table S5 Michaelis-Menten constant ($K_M$) values for third-generation cathepsin S substrates.** $K_M$ values were determined to assess the substrate binding affinity of cathepsin S for each third-generation peptide sequence. Measurements were carried out under steady-state conditions, and the data represent the average of at least two independent experiments.

| P4-P2 | P1 | Code | $K_M$, µM | | | |
|---|---|---|---|---|---|---|
| | | | Cathepsin S | Cathepsin B | Cathepsin L | Cathepsin V |
| Phe(F5)-Cit-2Aoc | Arg | **OG-64** | 8,53 | 8,52 | 10,9 | 16,5 |
| Phe(F5)-Cit-NptGly | Arg | **OG-191** | 62,2 | 62,2 | 169 | 59,9 |
| | Gln | **OG-192** | 21,2 | 20,4 | 63,7 | 27,1 |
| | Glu(Me) | **OG-193** | 15,2 | 12,6 | 41,0 | 22,5 |
| | Cit | **OG-194** | 9,43 | 24,9 | 72,9 | 22,1 |
| Phe(F5)-Cit-Lys(2ClZ) | Arg | **OG-195** | 4,39 | 5,33 | 19,5 | 18,3 |
| | Gln | **OG-196** | 0,27 | 0,88 | 23,0 | 13,4 |
| | Glu(Me) | **OG-197** | 0,74 | 3,63 | 42,8 | 19,4 |
| | Cit | **OG-198** | 0,62 | 2,67 | 36,5 | 12,8 |
| Phe(F5)-Thz-2Aoc | Arg | **OG-199** | 8,70 | 74,7 | 19,3 | 11,4 |
| | Gln | **OG-200** | 1,63 | 20,3 | 15,9 | 4,31 |
| | Glu(Me) | **OG-201** | 3,04 | 15,8 | 12,1 | 11,2 |
| | Cit | **OG-202** | 8,37 | 21,3 | 24,4 | 4,94 |
| Met(O2)-Cit-2Aoc | Arg | **OG-203** | 11,3 | 10,1 | 15,8 | 12,9 |
| | Gln | **OG-204** | 7,24 | 10,0 | 10,1 | 8,35 |
| | Glu(Me) | **OG-205** | 6,54 | 16,9 | 58,6 | 25,8 |
| | Cit | **OG-206** | 9,52 | 10,9 | 11,7 | 7,65 |
| Met(O2)-Cit-NptGly | Arg | **OG-207** | 42,5 | 201 | 133 | 118 |
| | Gln | **OG-208** | 48,1 | 294 | 105 | 102 |
| | Glu(Me) | **OG-209** | 39,2 | 369 | 236 | 176 |
| | Cit | **OG-210** | 32,9 | 322 | 242 | 54,4 |

**Table S6 $k_{cat}$ values for third-generation cathepsin S substrates.** $k_{cat}$ values were determined to assess the cathepsin S turnover toward 3rd generation fluorescent substrates. Measurements were carried out under steady-state conditions, and the data represent the average of at least two independent experiments.

| P4-P2 | P1 | Code | $k_{cat}$, $M^{-1}$ | | | |
|---|---|---|---|---|---|---|
| | | | Cathepsin S | Cathepsin B | Cathepsin L | Cathepsin V |
| Phe(F5)-Cit-2Aoc | Arg | **OG-64** | 4,90 | 1,23 | 0,15 | 0,42 |
| Phe(F5)-Cit-NptGly | Arg | **OG-191** | 23,76 | 2,00 | 0,85 | 5,56 |
| | Gln | **OG-192** | 2,52 | 0,13 | 0,39 | 0,76 |
| | Glu(Me) | **OG-193** | 2,02 | 0,10 | 0,32 | 0,51 |
| | Cit | **OG-194** | 1,08 | 0,28 | 0,49 | 0,85 |
| Phe(F5)-Cit-Lys(2ClZ) | Arg | **OG-195** | 1,60 | 0,39 | 0,20 | 0,26 |
| | Gln | **OG-196** | 0,19 | 0,04 | 0,12 | 0,14 |
| | Glu(Me) | **OG-197** | 0,26 | 0,02 | 0,09 | 0,15 |
| | Cit | **OG-198** | 0,27 | 0,05 | 0,16 | 0,23 |
| Phe(F5)-Thz-2Aoc | Arg | **OG-199** | 4,33 | 8,90 | 0,24 | 0,44 |
| | Gln | **OG-200** | 0,15 | 0,30 | 0,02 | 0,20 |
| | Glu(Me) | **OG-201** | 0,65 | 0,46 | 0,03 | 0,29 |
| | Cit | **OG-202** | 2,07 | 1,16 | 0,07 | 0,28 |
| Met(O2)-Cit-2Aoc | Arg | **OG-203** | 12,40 | 2,34 | 1,61 | 0,18 |
| | Gln | **OG-204** | 1,94 | 1,86 | 0,25 | 0,27 |
| | Glu(Me) | **OG-205** | 5,67 | 2,15 | 0,56 | 0,57 |
| | Cit | **OG-206** | 1,58 | 1,05 | 0,14 | 0,13 |
| Met(O2)-Cit-NptGly | Arg | **OG-207** | 22,48 | 4,40 | 6,74 | 9,89 |
| | Gln | **OG-208** | 4,40 | 1,09 | 0,94 | 3,74 |
| | Glu(Me) | **OG-209** | 22,11 | 1,88 | 1,96 | 5,83 |
| | Cit | **OG-210** | 7,21 | 2,41 | 2,37 | 2,02 |

**Table S7 Structures of cathepsin S-selective inhibitors** with the general formula Ac-Met($O_2$)-Cit-NptGly-P1-AOMK (JN1-JN6) or Ac-Phe($F_5$)-Cit-Lys(2ClZ)-P1-AOMK (JN7-JN12). Each inhibitor is based on a selective tetrapeptide scaffold and contains a reactive AOMK warhead at the C-terminus. Variations at the P1 position were introduced to fine-tune potency and selectivity toward cathepsin S.

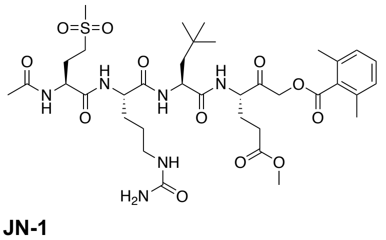

**JN-1**
Ac-Met(O2)-Cit-NptGly-Glu(Me)-AOMK

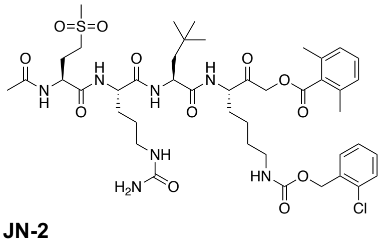

**JN-2**
Ac-Met(O2)-Cit-NptGly-Lys(2ClZ)-AOMK

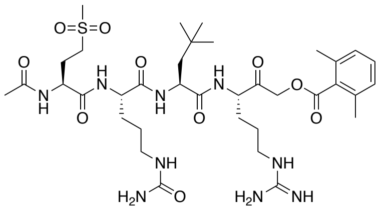

**JN-3**
Ac-Met(O2)-Cit-NptGly-Arg-AOMK

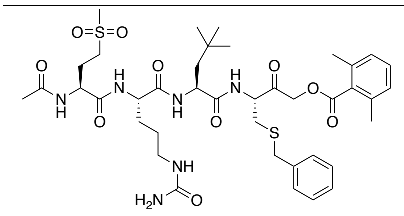

**JN-4**
Ac-Met(O2)-Cit-NptGly-Cys(Bzl)-AOMK

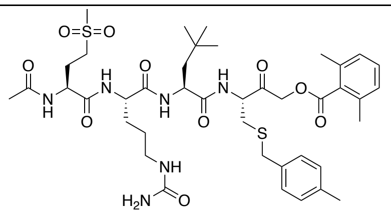

**JN-5**
Ac-Met(O2)-Cit-NptGly-Cys(MeBzl)-AOMK

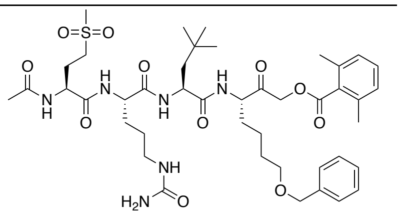

**JN-6**
Ac-Met(O2)-Cit-NptGly-Nle(OBzl)-AOMK

| | | |
|---|---|---|
| 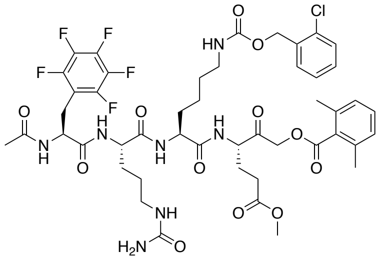<br>**JN-7**<br>Ac-Phe($F_5$)-Cit-Lys(2ClZ)-Glu(Me)-AOMK | 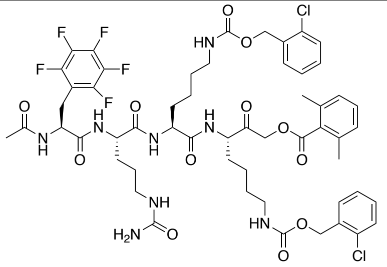<br>**JN-8**<br>Ac-Phe($F_5$)-Cit-Lys(2ClZ)-Lys(2ClZ)-AOMK | 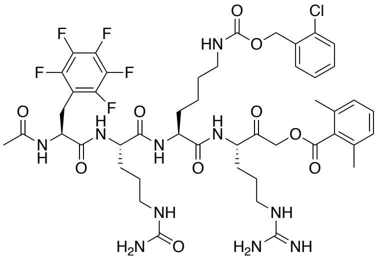<br>**JN-9**<br>Ac-Phe($F_5$)-Cit-Lys(2ClZ)-Arg-AOMK |
| 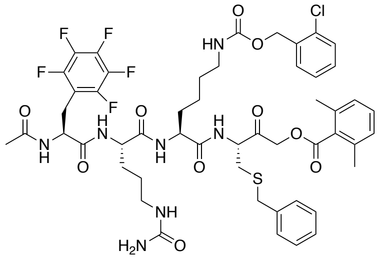<br>**JN-10**<br>Ac-Phe($F_5$)-Cit-Lys(2ClZ)-Cys(Bzl)-AOMK | 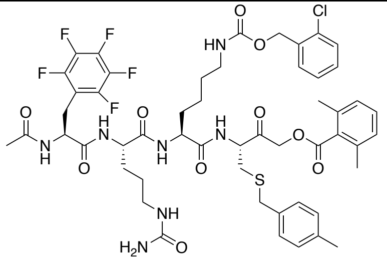<br>**JN-11**<br>Ac-Phe($F_5$)-Cit-Lys(2ClZ)-Cys(MeBzl)-AOMK | 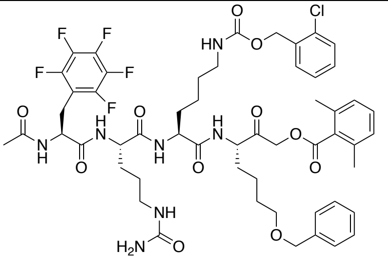<br>**JN-12**<br>Ac-Phe($F_5$)-Cit-Lys(2ClZ)-Nle(OBzl)-AOMK |

**Table S8 Second-order rate constants of cathepsin S inhibitors ($k_{obs}$/[I]).** Kinetic analysis of cathepsin S-selective inhibitors was performed to determine their second-order rate constants of inhibition ($k_{obs}$/[I]). Values were obtained under pseudo-first-order conditions by monitoring the time-dependent decrease in enzyme activity. Higher kobs/[I] values indicate more efficient inhibition.

| P4-P2 | P1 | Code | $k_{obs}$/[I], $M^{-1}s^{-1}$ | | |
|---|---|---|---|---|---|
| | | | Cathepsin S | Cathepsin L | Cathepsin B |
| Met($O_2$)-Cit-NptGly | Glu(Me) | **JN-1** | 392,000 | 560 | 7,440 |
| | Lys(2ClZ) | **JN-2** | 269,000 | 220 | 5,100 |
| | Arg | **JN-3** | 236,000 | 4,840 | 41,500 |
| | Cys(Bzl) | **JN-4** | 443,000 | 450 | 22,400 |
| | Cys(MeBzl) | **JN-5** | 344,000 | 610 | 10,500 |
| | Nle(OBzl) | **JN-6** | 246,000 | 650 | 16,900 |
| Phe($F_5$)-Cit-Lys(2ClZ) | Glu(Me) | **JN-7** | 609,999 | 870 | 17,000 |
| | Lys(2ClZ) | **JN-8** | 225,000 | 350 | 1,600 |
| | Arg | **JN-9** | 371,000 | 4,270 | 148,000 |
| | Cys(Bzl) | **JN-10** | 545,000 | 620 | 16,100 |
| | Cys(MeBzl) | **JN-11** | 261,000 | 320 | 1,900 |
| | Nle(OBzl) | **JN-12** | 392,000 | 490 | 5,160 |
| | | **E-64** | 132,000 | 65,100 | 81,200 |

**Table S9 Structures of cathepsin S-selective activity-based probes (ABPs) based on optimized substrate sequences.** Table presents the chemical structures of fluorescent ABPs designed using the most selective substrate motifs for cathepsin S. Each probe consists of a peptide recognition sequence, an irreversible AOMK warhead, and a fluorescent dye (Cy5 or BODIPY) conjugated to the N-terminus. The design preserves enzymatic selectivity while enabling probe visualization in biochemical and cellular assays.

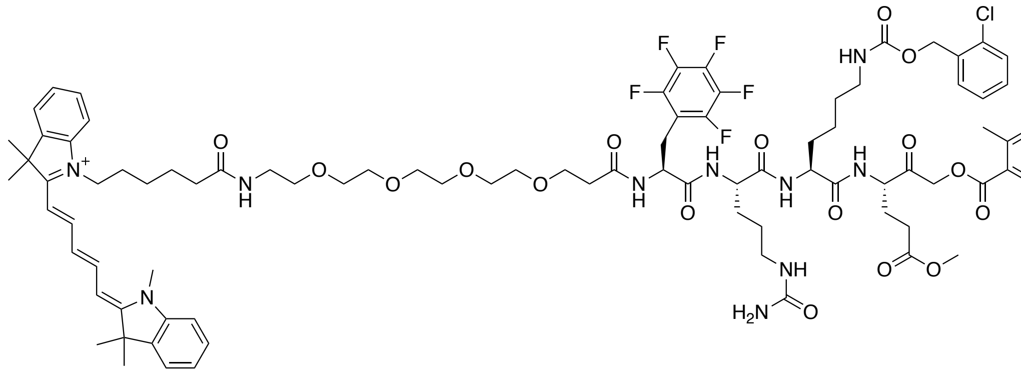

**OG-233**
Cy5-PEG(4)-Phe($F_5$)-Cit-Lys(2ClZ)-Glu(Me)-AOMK

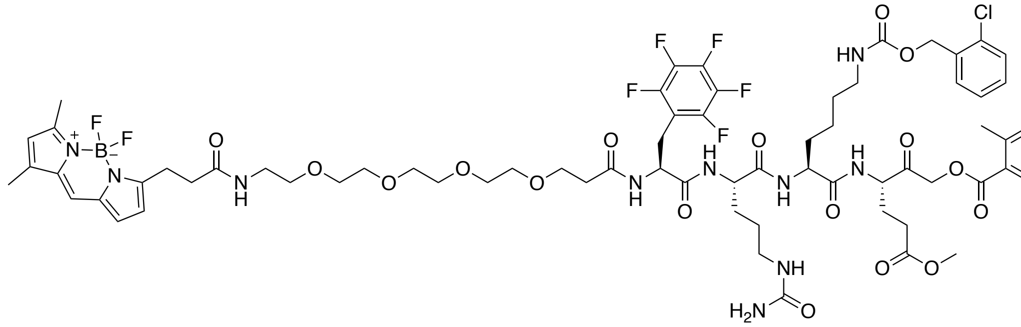

**OG-235**
BODIPY-PEG(4)-Phe($F_5$)-Cit-Lys(2ClZ)-Glu(Me)-AOMK

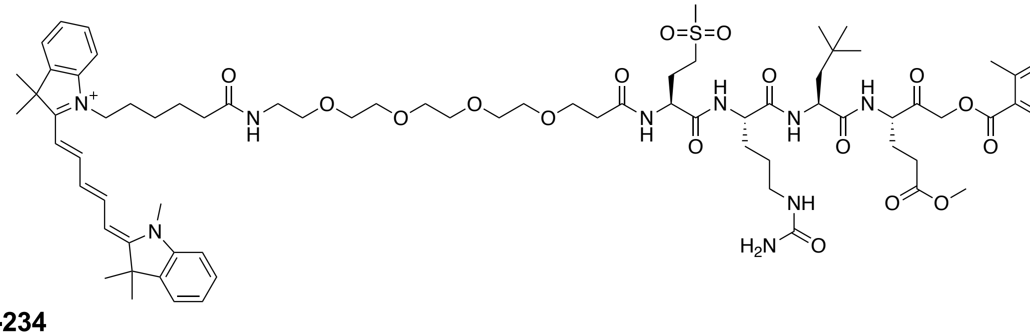

**OG-234**
Cy5-PEG(4)-Met($O_2$)-Cit-NptGly-Glu(Me)-AOMK

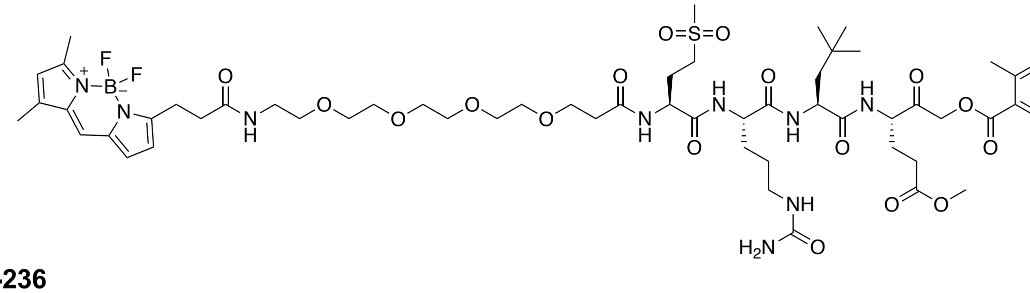

**OG-236**
BODIPY-PEG(4)-Met($O_2$)-Cit-NptGly-Glu(Me)-AOMK

**Table S10 Chemical structures of cathepsin S-selective peptide prodrugs** (OG-313, OG-314), non-cleavable control (OG-352), and pan-cathepsin prodrug (Z-Val-Cit-PABC-MMAE). This table displays the structures of MMAE-based prodrugs incorporating protease-cleavable peptide linkers. OG-313 and OG-314 contain cathepsin S-selective sequences identified through substrate optimization. OG-352 includes a non-cleavable D-amino acid at the P1 position, serving as a negative control. The widely used Z-Val-Cit-PABC-MMAE is included as a pan-cathepsin-cleavable reference.

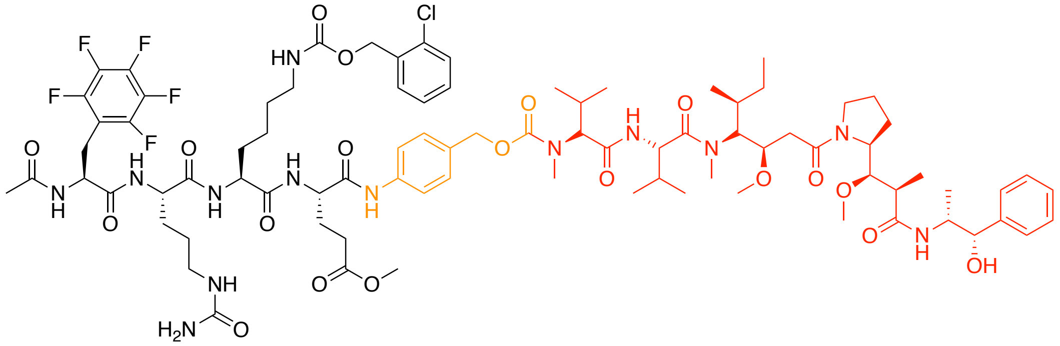

**OG-313**
Ac-Phe($F_5$)-Cit-Lys(2ClZ)-Glu(Me)-PABC-MMAE

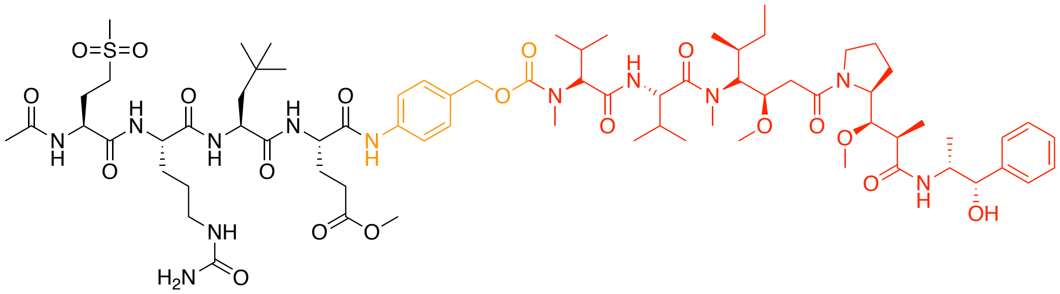

**OG-314**
Ac-Met($O_2$)-Cit-NptGly-Glu(Me)-PABC-MMAE

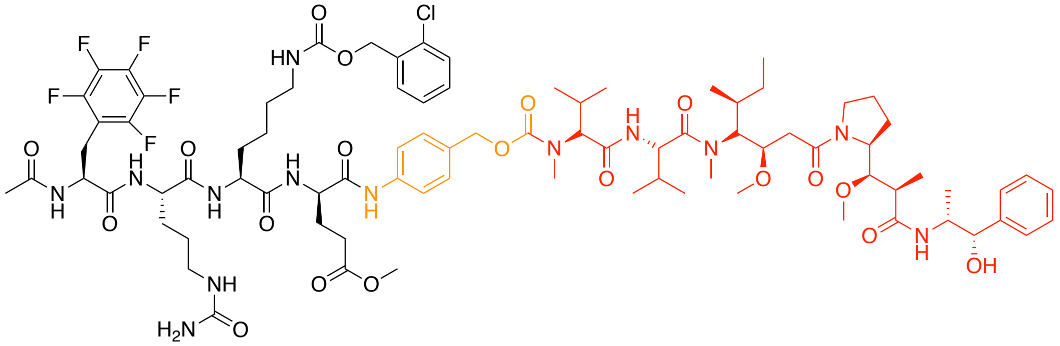

**OG-352**
Ac-Phe($F_5$)-Cit-Lys(2ClZ)-DGlu(Me)-PABC-MMAE

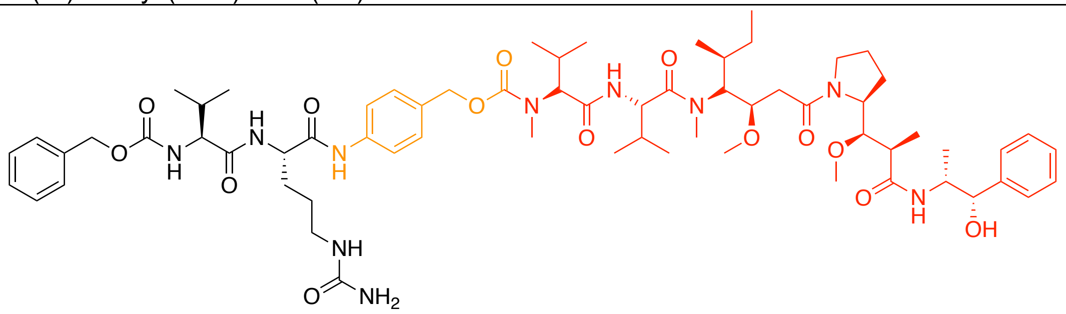

**Z-Val-Cit**
Cbz-Val-Cit-PABC-MMAE

**Table S11 Metal-conjugated antibody panel for mass cytometry profiling of breast tumor architecture.** This panel includes antibodies targeting markers for major cell populations within the tumor microenvironment, including apoptotic cells, immune cells, epithelial cells, fibroblasts, and endothelial cells. The table also outlines the gating strategies used to define each cell type, providing a framework for high-resolution single-cell identification and population profiling by CyTOF.

| Cell type | antibody | metal |
|---|---|---|
| Apoptotic cells (**cPARP+, cCasp3+)** | Anti-cPARP | 143Nd |
| | Anti-cl. Casp-3 | 142Ce |
| Granulocytes (**CD66b+**) | Anti-CD66b | 113Cd |
| Lymphocytes (**CD45+**, CD66b-) | Anti-CD45 | 89Y |
| B cells (CD45+, **CD19+**) | Anti-CD19 | 165Ho |
| T cells (CD45+, **CD3+**) | Anti-CD3 | 154Sm |
| CD4+ T cells (CD45+, CD3+, **CD4+**, CD8-) | Anti-CD4 | 110Cd |
| CD8+ T cells (CD45+, CD3+, CD4-, **CD8+**) | Anti-CD8 | 114Cd |
| Macrophages (CD45+, CD3-, CD19-, **CD11b+**) | Anti-CD11b | 209Bi |
| NK cells (CD45+, CD3-, CD19-, **CD56+**) | Anti-CD56 | 176Yb |
| Other lymphocytes (CD45+, CD3-, CD19-, CD11b-, CD56-) | - | - |
| Epithelial cells (CD45-, CD66b-, **EpCAM+** or **Cadh3+**) | Anti-EpCAM | 141Pr |
| | Anti-Cadherin-3 | 172Yb |
| Fibroblasts (CD45-, CD66b-, EpCAM-, Cadh3-, **FAP+ or SMA+**) | Anti-FAP | 164Dy |
| | Anti-SMA | 161Dy |
| Endothelial (CD45-, CD66b-, EpCAM-, Cadh3-, FAP+, SMA-, **CD31+**) | Anti-CD31 | 144Nd |
| Other non-immune (all markers negative) | - | - |

**Table S12 Metal-conjugated antibody panel for detection of tumor-associated biomarkers and cathepsin S by mass cytometry.** This panel includes antibodies specific for key breast cancer markers such as HER2, estrogen receptor (ER), and progesterone receptor (PR), enabling molecular subtyping of tumor cells. Additionally, the panel features a metal-conjugated antibody against cathepsin S, allowing single-cell resolution profiling of protease expression within the tumor microenvironment.

| Protein/hormone target | antibody | metal |
|---|---|---|
| HER-2 | Anti-HER-2 | 113Cd |
| Progesterone (PR) | Anti-PR | 145Nd |
| Estrogen (ER) | Anti-ER | 163Dy |
| Cathepsin S | Anti-Cat S | 155Gd |
| Cathepsin B | Anti-Cat B | 162Dy |

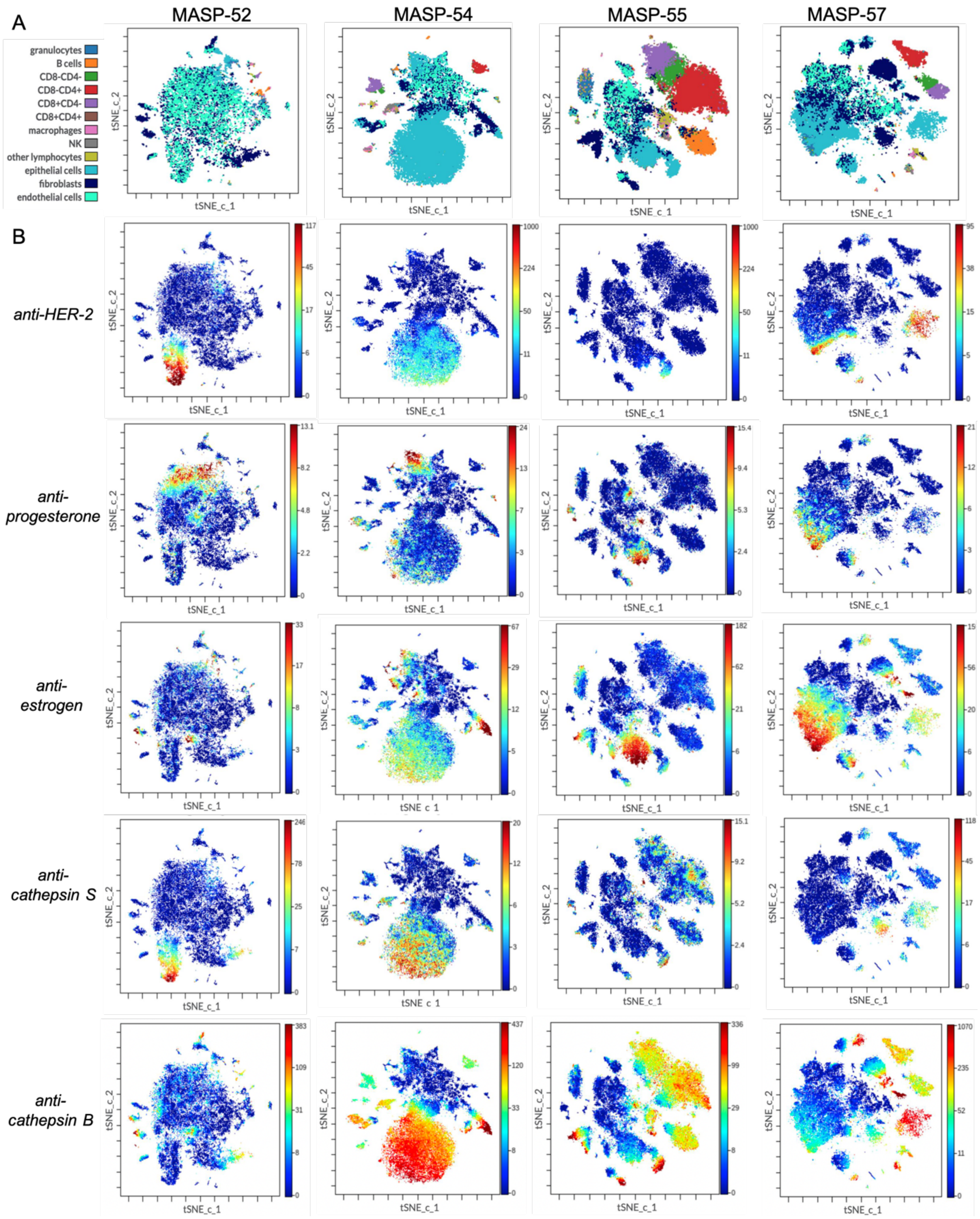


**Figure S4 Single-cell mass cytometry profiling of the breast tumor microenvironment.** Freshly resected breast tumors were enzymatically dissociated into single-cell suspensions and analyzed by mass cytometry (CyTOF) to generate high-dimensional cellular landscapes. **(A)** viSNE analysis revealed marked intratumoral heterogeneity among patient samples. While epithelial and endothelial cells represented the dominant populations across all tumors, the abundance of tumor-infiltrating immune cells varied significantly, with some patients exhibiting substantial immune infiltration. **(B)** viSNE-based marker distribution maps showed high co-expression of HER2 with cathepsin S, whereas correlations with PR and ER were only moderate. HER2 expression was primarily restricted to epithelial cells, while cathepsin S was detected in both epithelial cells and infiltrating immune cells. These findings support the rationale for developing protease-selective ADCs that exploit cathepsin S activity in HER2-positive tumors or enable extracellular activation in immune cell-rich tumor microenvironments.

**Table S13** The LC-MS analysis of substrates, inhibitors, activtiy-based probes and prodrugs for cathepsin S *(to be completed)*

| Code | Compound structure | $[M+H]^+$ calculated | $[M+H]^+$ measured |
|---|---|---|---|
| **OG-64** | Ac-Phe($F_5$)-Cit-2Aoc-Arg-ACC | 952,41 | 952,49 |
| **OG-65** | Ac-Phe($F_5$)-Idc-2Aoc-Arg-ACC | 940,38 | 940,45 |
| **OG-66** | Ac-Phe($F_5$)-Thz-Lys(2ClZ)-Arg-ACC | 1065,31 | 1065,45 |
| **OG-67** | Ac-Phe($F_5$)-Cit-3Pal-Arg-ACC | 958,37 | 958,41 |
| **OG-68** | Ac-Phe($F_5$)-Tic-2Aoc-Arg-ACC | 954,39 | 954,44 |
| **OG-69** | Ac-Phe($F_5$)-Arg-Lys(2ClZ)-Arg-ACC | 1106,40 | 1106,56 |
| **OG-70** | HN-Idc-Cit-Leu-Arg-ACC | 788,38 | 788,47 |
| **OG-71** | HN-Idc-Thz-Leu-Arg-ACC | 746,30 | 746,55 |
| **OG-72** | HN-Idc-Ile-3Pal-Arg-ACC | 779,36 | 779,45 |
| **OG-73** | HN-Idc-Tic-2Aoc-Arg-ACC | 818,40 | 818,50 |
| **OG-74** | HN-Idc-Arg-Leu-Arg-ACC | 787,39 | 787,48 |
| **OG-75** | Ac-Pro-Arg-Leu-Arg-ACC | 783,42 | 783,50 |
| **OG-76** | Ac-Pro-Ile-3Pal-Arg-ACC | 775,38 | 775,45 |
| **OG-77** | Ac-Pro-Thz-Lys(2ClZ)-Arg-ACC | 925,34 | 925,39 |
| **OG-79** | Ac-Pro-Arg-2Aoc-Arg-ACC | 811,45 | 811,53 |
| **OG-80** | Ac-Pro-Arg-Hyp(Bzl)-Arg-ACC | 873,43 | 873,52 |
| **OG-81** | Ac-Tle-Cit-2Aoc-Arg-ACC | 828,47 | 828,55 |
| **OG-82** | HN-Idc-Lys(2ClZ)-Arg-ACC | 814,30 | 814,36 |
| **OG-83** | Ac-Tle-Thz-2Aoc-Arg-ACC | 786,39 | 786,46 |
| **OG-84** | Ac-Tle-Ile-3Pal-Arg-ACC | 791,41 | 791,50 |
| **OG-85** | Ac-Tle-Arg-Hyp(Bzl)-Arg-ACC | 889,46 | 889,53 |
| **OG-86** | Ac-Tle-Arg-2Aoc-Arg-ACC | 827,48 | 827,58 |
| **OG-87** | Ac-Asp(Me)-Ile-3Pal-Arg-ACC | 807,37 | 807,47 |
| **OG-88** | Ac-Asp(Me)-Tic-Leu-Arg-ACC | 818,38 | 818,46 |
| **OG-89** | Ac-Asp-Tic-Hyp(Bzl)-Arg-ACC | 894,37 | 894,42 |
| **OG-90** | Ac-Asp(Me)-Thz-Lys(2ClZ)-Arg-ACC | 957,33 | 957,38 |
| **OG-91** | Ac-Met($O_2$)-Cit-Hyp(Bzl)-Arg-ACC | 940,39 | 940,45 |
| **OG-93** | Ac-Met($O_2$)-Thz-Lys(2ClZ)-Arg-ACC | 991,31 | 991,56 |
| **OG-94** | Ac-Met($O_2$)-Ile-Hyp(Bzl)-Arg-ACC | 896,39 | 896,47 |
| **OG-95** | Ac-Bpa-Tic-Hyp(Bzl)-Arg-ACC | 1030,44 | 1030,52 |
| **OG-96** | Ac-Bpa-Arg-2Aoc-Arg-ACC | 965,49 | 965,54 |

| | | | |
|---|---|---|---|
| **OG-115** | Ac-1Nal-Cit-2Aoc-Arg-ACC | 912,47 | 912,55 |
| **OG-116** | Ac-Bpa-Cit-2Aoc-Arg-ACC | 966,48 | 966,58 |
| **OG-117** | Ac-Met($O_2$)-Cit-2Aoc-Arg-ACC | 878,41 | 878,50 |
| **OG-118** | Ac-Leu-Cit-2Aoc-Arg-ACC | 828,47 | 828,57 |
| **OG-119** | Ac-Phe($F_5$)-Cit-Leu-Arg-ACC | 924,37 | 924,46 |
| **OG-120** | Ac-Phe($F_5$)-Cit-Nle-Arg-ACC | 924,37 | 924,48 |
| **OG-121** | Ac-Leu-Thz-Lys(2ClZ)-Arg-ACC | 941,37 | 941,45 |
| **OG-122** | Ac-Leu-Thz-hLeu-Arg-ACC | 772,37 | 772,47 |
| **OG-123** | Ac-Leu-Thz-NptGly-Arg-ACC | 772,37 | 772,49 |
| **OG-133** | Ac-Phe($F_5$)-Cit-2Aoc-Gln-ACC | 924,36 | 924,44 |
| **OG-134** | Ac-Phe($F_5$)-Cit-2Aoc-Nle-ACC | 909,39 | 909,49 |
| **OG-135** | Ac-Phe($F_5$)-Cit-2Aoc-Glu(Me)-ACC | 939,36 | 939,44 |
| **OG-136** | Ac-Phe($F_5$)-Cit-2Aoc-Cit-ACC | 953,39 | 953,49 |
| **OG-137** | Ac-Phe($F_5$)-Cit-2Aoc-Lys(TFA)-ACC | 1020,38 | 1020,51 |
| **OG-138** | Ac-Phe($F_5$)-Cit-2Aoc-Thr(Bzl)-ACC | 987,40 | 987,44 |
| **OG-139** | Ac-Phe($F_5$)-Cit-2Aoc-Cys(MeOBzl)-ACC | 1019,37 | 1019,51 |
| **OG-140** | Ac-Phe($F_5$)-Cit-2Aoc-Cys(MeBzl)-ACC | 1003,37 | 1003,50 |
| **OG-141** | Ac-Phe($F_5$)-Cit-2Aoc-hTyr(Me)-ACC | 987,40 | 987,45 |
| **OG-142** | Ac-Phe($F_5$)-Cit-2Aoc-Nle(OBzl)-ACC | 1015,43 | 1015,53 |
| **OG-143** | Ac-Phe($F_5$)-Cit-2Aoc-Met-ACC | 927,34 | 927,43 |
| **OG-144** | Ac-Phe($F_5$)-Cit-2Aoc-Abu-ACC | 881,35 | 881,42 |
| **OG-145** | Ac-Phe($F_5$)-Cit-2Aoc-Nva-ACC | 895,37 | 895,46 |
| **OG-146** | Ac-Phe($F_5$)-Cit-2Aoc-hLeu-ACC | 923,40 | 923,50 |
| **OG-147** | Ac-Phe($F_5$)-Cit-2Aoc-2Aoc-ACC | 937,42 | 937,49 |
| **OG-191** | Ac-Phe($F_5$)-Cit-NptGly-Arg-ACC | 938,39 | 938,45 |
| **OG-192** | Ac-Phe($F_5$)-Cit-NptGly-Gln-ACC | 910,34 | 910,41 |
| **OG-193** | Ac-Phe($F_5$)-Cit-NptGly-Glu(Me)-ACC | 925,34 | 925,42 |
| **OG-194** | Ac-Phe($F_5$)-Cit-NptGly-Cit-ACC | 939,37 | 939,42 |
| **OG-195** | Ac-Phe($F_5$)-Cit-Lys(2ClZ)-Arg-ACC | 1107,38 | 1107,54 |
| **OG-196** | Ac-Phe($F_5$)-Cit-Lys(2ClZ)-Gln-ACC | 1079,34 | 1079,45 |
| **OG-197** | Ac-Phe($F_5$)-Cit-Lys(2ClZ)-Glu(Me)-ACC | 1094,34 | 1094,48 |
| **OG-198** | Ac-Phe($F_5$)-Cit-Lys(2ClZ)-Cit-ACC | 1108,36 | 1108,51 |
| **OG-199** | Ac-Phe($F_5$)-Thz-2Aoc-Arg-ACC | 910,33 | 910,40 |
| **OG-200** | Ac-Phe($F_5$)-Thz-2Aoc-Glu-ACC | 883,27 | 883,30 |

| | | | |
|---|---|---|---|
| **OG-201** | Ac-Phe($F_5$)-Thz-2Aoc-Gln-ACC | 882,28 | 882,34 |
| **OG-202** | Ac-Phe($F_5$)-Thz-2Aoc-Cit-ACC | 911,31 | 911,37 |
| **OG-203** | Ac-Met($O_2$)-Cit-2Aoc-Arg-ACC | 878,41 | 878,49 |
| **OG-204** | Ac-Met($O_2$)-Cit-2Aoc-Gln-ACC | 850,37 | 850,43 |
| **OG-205** | Ac-Met($O_2$)-Cit-2Aoc-Glu(Me)-ACC | 865,37 | 865,44 |
| **OG-206** | Ac-Met($O_2$)-Cit-2Aoc-Cit-ACC | 879,40 | 879,46 |
| **OG-207** | Ac-Met($O_2$)-Cit-NptGly-Arg-ACC | 864,40 | 864,49 |
| **OG-208** | Ac-Met($O_2$)-Cit-NptGly-GlnA-CC | 836,35 | 836,43 |
| **OG-209** | Ac-Met($O_2$)-Cit-NptGly-Glu(Me)-ACC | 851,35 | 851,42 |
| **OG-210** | Ac-Met($O_2$)-Cit-NptGly-Cit-ACC | 865,35 | 865,48 |
| **JN-1** | Ac-Met($O_2$)-Cit-NptGly-Glu(Me)-AOMK | 797,37 | 797,45 |
| **JN-2** | Ac-Met($O_2$)-Cit-NptGly-Lys(2ClZ)-AOMK | 950,40 | 950,41 |
| **JN-3** | Ac-Met($O_2$)-Cit-NptGly-Arg-AOMK | 810,41 | 810,43 |
| **JN-4** | Ac-Met($O_2$)-Cit-NptGly-Cys(Bzl)-AOMK | 847,37 | 847,47 |
| **JN-5** | Ac-Met($O_2$)-Cit-NptGly-Cys(MeBzl)-AOMK | 861,38 | 861,49 |
| **JN-6** | Ac-Met($O_2$)-Cit-NptGly-Nle(OBzl)-AOMK | 873,44 | 873,52 |
| **JN-7** | Ac-Phe($F_5$)-Cit-Lys(2ClZ)-Glu(Me)-AOMK | 1040,35 | 1040,40 |
| **JN-8** | Ac-Phe($F_5$)-Cit-Lys(2ClZ)-Lys(2ClZ)-AOMK | 1193,39 | 1193,54 |
| **JN-9** | Ac-Phe($F_5$)-Cit-Lys(2ClZ)-Arg-AOMK | 1053,39 | 1054,00 |
| **JN-10** | Ac-Phe($F_5$)-Cit-Lys(2ClZ)-Cys(Bzl)-AOMK | 1090,35 | 1090,52 |
| **JN-11** | Ac-Phe($F_5$)-Cit-Lys(2ClZ)-Cys(MeBzl)-AOMK | 1104,37 | 1104,55 |
| **JN-12** | Ac-Phe($F_5$)-Cit-Lys(2ClZ)-Nle(OBzl)-AOMK | 1116,42 | 1116,48 |
| **OG-233** | Cy5-PEG(4)-Phe($F_5$)-Cit-Lys(2ClZ)-Glu(Me)-AOMK | 1709,7<br>855,39 $[M+H]^{2+}$ | 855,76 $[M+H]^{2+}$ |
| **OG-235** | BODIPY-PEG(4)-Phe($F_5$)-Cit-Lys(2ClZ)-Glu(Me)-AOMK | 1518,59<br>760,30 $[M+H]^{2+}$ | 760,58 $[M+H]^{2+}$ |
| **OG-234** | Cy5-PEG(4)-Met($O_2$)-Cit-NptGly-Glu(Me)-AOMK | 1466,79<br>733,90 $[M+H]^{2+}$ | 733,35 $[M+H]^{2+}$ |
| **OG-236** | BODIPY-PEG(4)-Met($O_2$)-Cit-NptGly-Glu(Me)-AOMK | 1275,61<br>638,81 $[M+H]^{2+}$ | 638,66 $[M+H]^{2+}$ |
| **OG-313** | Ac-Phe($F_5$)-Cit-Lys(2ClZ)-Glu(Me)-PABC-MMAE | 1742,83<br>871,92 $[M+H]^{2+}$ | 872,18 $[M+H]^{2+}$ |
| **OG-314** | Ac-Met($O_2$)-Cit-NptGly-Glu(Me)-PABC-MMAE | 1499,84<br>750,42 $[M+H]^{2+}$ | 750,80 $[M+H]^{2+}$ |
| **OG-352** | Ac-Phe($F_5$)-Cit-Lys(2ClZ)-DGlu(Me)-PABC-MMAE | 1742,83<br>871,92 $[M+H]^{2+}$ | 872,12 $[M+H]^{2+}$ |
| **Z-Val-Cit** | Cbz-Val-Cit-PABC-MMAE | 1257,74<br>629,38 $[M+H]^{2+}$ | 629,63 $[M+H]^{2+}$ |
